# PREMISE: A Quality-Aware Probabilistic Framework for High-Resolution Source Assignment in Metagenomic Sequencing

**DOI:** 10.64898/2026.03.15.711921

**Authors:** Sriram Vijendran, Karin Dorman, Tavis K. Anderson, Oliver Eulenstein

**Affiliations:** Iowa State University, Ames IA 50014, USA; National Animal Disease Center, Agricultural Research Service, United States Department of Agriculture, Ames IA 50010, USA

**Keywords:** Metagenomics, Taxonomic Classification, Expectation Maximization, Rust, Next Generation Sequencing (NGS)

## Abstract

Metagenomic applications, from influenza A virus (IAV) subtype tracking to bacterial strain resolution, frequently require sub-species resolution. However, traditional *k*-mer-based methods fragment reads into short *k*-mers and discard per-base quality scores. By omitting both intra-read linkage and error information, these methods confound closely related genomic variants, thereby impeding real-time pathogen surveillance and containment. We introduce PREMISE (Pathogen Resolution via Expectation-Maximization In Sequencing Experiments), a probabilistic, alignment-based framework implemented in Rust for high-resolution source assignment against near-complete reference databases. PREMISE aligns whole reads using a bidirectional run-length FM-index and quantifies reference abundances with a quality-aware Expectation-Maximization algorithm mapped over a taxonomy. This frame-work applies an optional sparsity penalty to resolve ambiguous assignments, which is particularly important when sparse samples are mapped against reference databases rich in highly homologous candidate references. On ten dense CAMI II strain-madness samples, PREMISE achieves the highest species-level read classification F1 score (0.992) among eight evaluated methods, with no false-positive or missed species. At the strain level, PREMISE maps reads to their source genome with top precision (0.80 vs. at most 0.64 for competing methods), achieves 100% recall, and estimates abundances as accurately as the best competing tool. These results demonstrate PREMISE’s ability to provide rigorous, uncertainty-aware source resolution across both dense and sparse metagenomes, accurately resolving targets ranging from viral subtypes to bacterial strains.

**Availability:** https://github.com/sriram98v/premise under an MIT license.

## 1 Introduction

High-throughput sequencing increasingly demands for strain-resolved answers, from tracking variants of viral pathogens to separating closely related strains in microbial communities [32]. Genomic surveillance with high throughput sequencing has transformed viral threat detection, monitoring, and management [26,12,54]: untargeted metagenomic sequencing can surveil the environment for emerging pathogens and reveal the etiology of idiopathic infections [54], and targeted whole genome sequencing shapes the public health management of SARS-CoV-2 [26,55] and influenza A [16,57]. Standardized enrichment protocols [65,21] enable the routine monitoring of clinical samples, perhaps pooled for efficiency [39], and of environmental samples such as wastewater [51], to detect novel variants and follow strain trajectories over time [50,53].

The main bioinformatic tasks associated with analyzing sequencing reads are (1) *high resolution* taxonomic classification of reads, (2) estimation of taxonomic abundances (profiling), and (3) identification of novel taxa. Classification and profiling remain computational bottlenecks [26], often with poor accuracy at the strain- or variant-level of interest in enriched samples [63,52], and misclassifications reduce diagnostic test validity and delay public health interventions [11]. High resolution requires comprehensive databases and valid statistical models for within-taxon genetic variation and residual read variation. Ongoing discovery [32] and computational limits [4] leave most databases incomplete. However, some viral disease databases are *near* complete and routinely updated [13,31], and the genomes of a defined, e.g. synthetic, community can be fully known. Paired with a valid model for library preparation and sequencing errors, such a database makes unprecedented resolution possible.

Two regimes arise against a near-complete database. In *dense* samples, most references in the database are present, as in the strain-madness samples of the second Critical Assessment of Metagenome Interpretation (CAMI II) [32], where hundreds of genomes from a few dozen taxa co-occur and many strains of one species share most of their sequence. In *sparse* samples, a few references are present among many highly homologous ones, as in an enriched influenza sample, in which the 8 segments of one or a few viruses are sampled from a database of thousands of segments. Dense samples demand that reads be divided among near-identical strains that are all present, whereas sparse samples demand that near-identical references that are absent be rejected. A method serving both must resolve strains where the reads support them and must otherwise report the taxon at which they become indistinguishable.

### Related Works

Profiling against complete databases was first attempted in RNA-seq, where RSEM [24], Kallisto [5], and Salmon [40,35] estimate transcript abundances by EM, and Metakallisto [46] and Salmon’s `--meta` option carry this quantification to metagenomes. Metagenomic tools that pose a probability model, such as Sigma [1], Karp [45], Mora [64], and Sylph [47], make simplifying assumptions—assuming iid sub-stitutions, ignoring indels, relying on the scores of heuristic aligners, or working on *k*-mer sketches—to cope with incomplete databases or to maintain competitive runtimes. Most metagenomic tools instead achieve validity through thresholds and filters [63] and trade resolution for speed. *k*-mer methods such as Kraken [60,59], KrakenUniq [6], CLARK [38,37], Bracken [27], Ganon [43,44], and KMCP [48] are fast.

In contrast, MEGAN [18], Centrifuge [19], Metalign [22], CAMMiQ [66], and PanTax [62] use partial or full read linkage, often with heavy filtering of candidate references. Benchmarks consistently find that false positives remain a serious problem and that performance drops sharply at fine resolution [61,10,32,52,33]. Supplementary Section S1 reviews these methods in detail.

### Our Contribution

We propose PREMISE (Pathogen Resolution via EM In Sequencing Experiments), a probabilistic framework implemented in Rust that combines a near-complete database with a sound model of read variation, bridging fast heuristic alignment with rigorous alignment needed for estimation. Using a bidirectional run-length FM-index (B-RLFM) [14,28,20,2], PREMISE seeds alignments with Super-Maximal Exact Matches and extends them without indels to compute read likelihoods from base-level Phred qualities.

References correspond to the leaves of a taxonomy, and EM on the taxonomy independently updates the conditional proportions for the children of each node. For sparse samples, an optional penalty promotes sparsity relative to the database and removes taxa not supported by the reads; for dense samples, this penalty is disabled. Each read is placed at the deepest node whose posterior probability exceeds a user-defined confidence threshold, so reads that the references cannot resolve stay at the species or genus level instead of being forced onto a particular strain. PREMISE is designed for well-characterized sources sequenced using low-indel technologies, and it serves as a basis for efficient approaches with sound statistical models that improve the reliability of metagenomic classifications and abundance profiles.

The remainder of this article is organized as follows. Section 2 describes the probabilistic model for read generation, the EM algorithm over the taxonomy, the seeding and alignment of reads to references, and the assignment of reads to taxa. Section 3 describes the dense CAMI II and sparse influenza samples and the evaluation metrics. Section 4 compares PREMISE with Centrifuger, Ganon, Karp, KMCP, Mora, Salmon, and Sylph, first on the dense and then on the sparse samples, and summarizes index construction and the sensitivity of PREMISE to its parameters. Section 5 discusses the results and future directions. The Supplementary Material gives the full review of related work, the derivations and implementation details of the methods, the dataset details, and the extended results.

## 2 Methods

We describe the read model, the EM algorithm that estimates the reference proportions over the taxonomy, its implementation, and the assignment of reads to taxa.

### 2.1 Preliminaries

Random variables (r.v.’s) are written in capital letters and their realizations in lowercase. Vectors are boldface and indexed from one, so ***v***[*j*] is the *j*^th^ element of ***v***; sequences (strings) are vectors over a finite alphabet *Σ*. We write |·| for the size of a set or the length of a vector, ⟬*i*⟭= {1, …, *i*},1(*b*) for the indicator of a boolean expression *b*, and 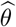 for an estimate of a parameter *θ*.

Let *T* be a rooted taxonomy with vertex set *V*, leaf set *L*, and root *r* (distinct from boldfaced reads ***r***_*i*_). For *v* ∈*V*, *ch*(*v*) denotes its children, *rTv* the root-to-*v* path, and *L*_*v*_ the leaves below *v*. Each of the *m* reference sequences ***D***_*l*_, distinct genomes, contigs, or segments, is attached as a leaf child of the most specific vertex it resolves to (usually species, but occasionally higher taxon; Figure 1a–b). Leaves and reference sequences are therefore in one-to-one correspondence: =*D* {***D***_*l*_} _*l∈L*_, |*L*| = *m*, and *L*_*v*_ indexes the set of references below *v*. When no taxonomy is available, every reference is a child of the root *r* in a star-like tree.

**Fig. 1:**
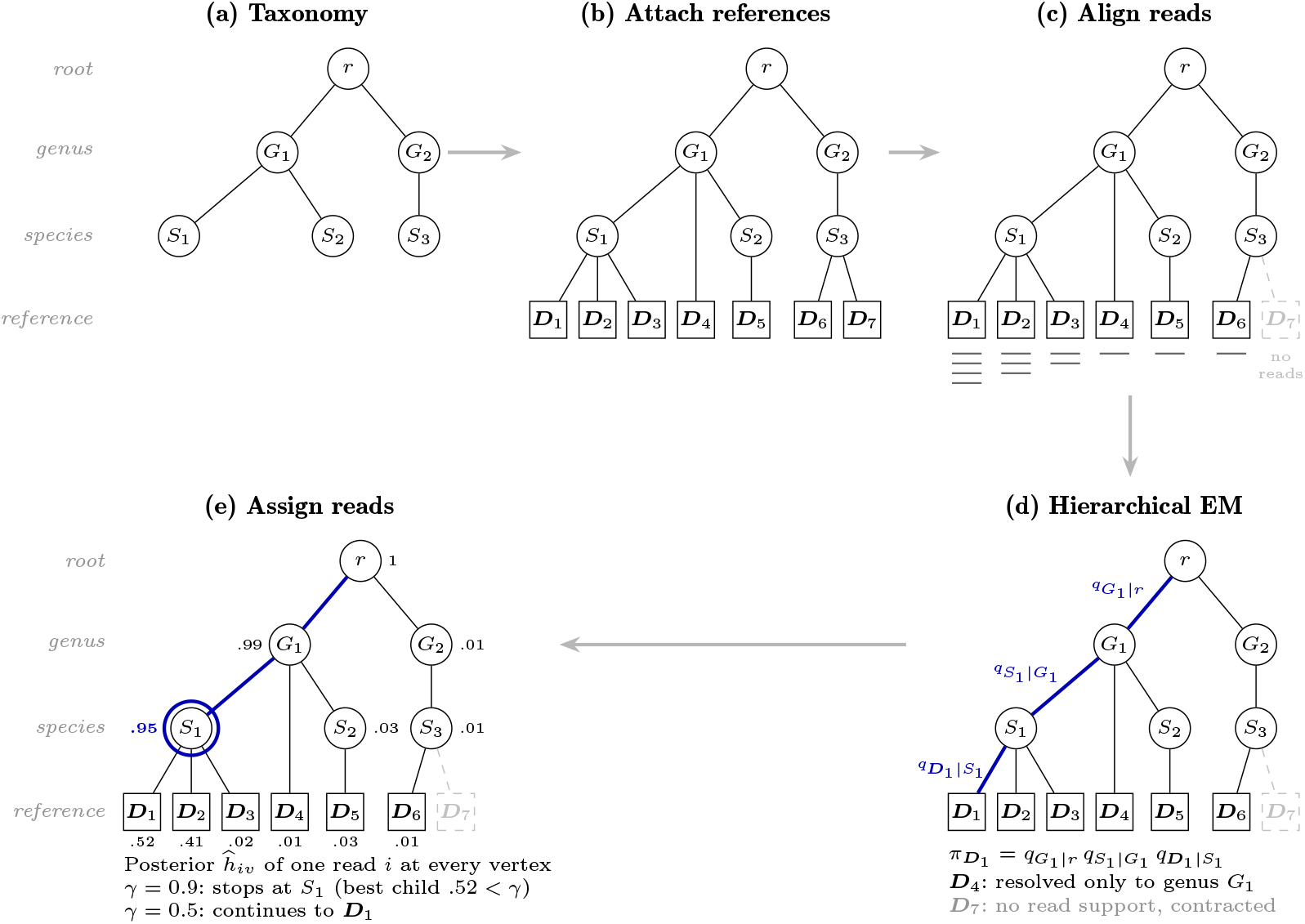
Taxonomy and read assignment in PREMISE. (a) A taxonomy. (b) Each reference ***D***_*l*_ (square) is resolved as leaf below its lowest assigned taxon. (c) Reads (bars) are aligned to references; without read support, ***D***_7_ is dropped. (d) EM estimates a conditional proportion *q*_*c*|*v*_ on every edge; proportion assigned to a vertex is the product of *q*_*c*|*v*_ along its root path. (e) After EM, the posterior probability of a read *i* assigned to an internal vertex *v* is the sum of the posterior assignment probabilities of it’s subtrees. From root, read *i* descends to most probable child while *ĥ*_*iv*_ *> γ*: setting *γ* = 0.9 assigns at species *S*_1_; with *γ* = 0.5, assignment is to ***D***_1_.

### 2.2 Read Generating Model

The observed data are a set of sequenced reads 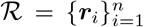 and their quality scores. We assume each read is an independent, noisy observation of one contiguous interval of a reference in *D*. We further assume that all technical sequences, such as adapters, primers, and barcodes, have been removed from the reads, that there are no insertion/deletion (indel) sequencing errors [34,36], and that the database is complete, i.e., it contains the source of every read. Each leaf *l* ∈*L* has a mixing proportion *π*_*l*_*≥*0, with *∑* _*l∈L*_ *π*_*l*_ = 1 that is to be estimated. We assign to each internal vertex *v* the total proportion of the leaves beneath it, 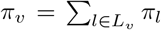, so that *π*_*r*_ = 1 and *π*_*v*_ = ∑_*c∈ch*(*v*)_ *π*_*c*_. The reads are sampled from a subset of references *S*= {*l* ∈*L* : *π*_*l*_ *>* 0} present in a population of interest. Each read *i* is generated in three steps. First, a leaf is drawn, *S*_*i*_ = *l* with probability *π*_*l*_. Second, since Illumina reads are much shorter than references, a start position *Z*_*i*_ ∈ [⟬|***D***_*l*_ | − |***r***_*i*_ | + 1⟭ is drawn with probability *η*_*ilz*_ := ℙ(*Z*_*i*_ = *z* | *S*_*i*_ = *l*). Third, the read is copied from ***D***_*l*_[*z*], …, ***D***_*l*_[*z* + |***r***_*i*_| − 1] with independent substitution errors at each site.

#### Read-level likelihood

Given *S*_*i*_ = *l* and *Z*_*i*_ = *z*, the probability of read ***r***_*i*_, *a*_*ilz*_ := P(***R***_*i*_ = ***r***_*i*_ *S*_*i*_ = *l, Z*_*i*_ = *z*), is

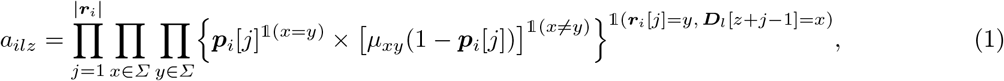

where ***p***_*i*_[*j*] ∈ [0, 1) is the probability that the *j*^th^ base call of the read matches the reference base ***D***_*l*_[*z* +*j* −1], and *µ*_*xy*_ = P(***R***_*i*_[*j*] = *y* | ***D***_*l*_[*z* + *j* − 1] = *x, S*_*i*_ = *l, Z*_*i*_ = *z*), with *µ*_*xx*_ = 0, *µ*_*xy*_ ≥ 0, and ∑_*y∈Σ*_ *µ*_*xy*_ = 1, is the probability of reading base *y* when an error has occurred and the true base is *x* ≠ *y*. Marginalizing over the start position gives conditional read likelihood

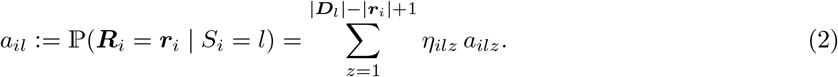

which is used during EM on the taxonomy (Section 2.4). There is insufficient replication to estimate ***p***_*i*_ as posed, so we model it with the Phred error model of the base call quality scores [8,41] available in FASTQ files, set *µ*_*xy*_ = 1*/*3 for *x* ≠*y*, and assume uniform coverage, *η*_*ilz*_ = 1*/*(|***D***_*l*_| − |***r***_*i*_| + 1).

### 2.3 Optimizing Read Classification

We estimate the proportions ***π*** with the EM algorithm, parameterized over the taxonomy (Figure 1d), optionally adding a sparsity-inducing penalty to the log-likelihood. For sparse samples |*S*| ≪ *m*, the penalty removes references with little read support; for dense samples, it is omitted (*ρ* = 0 below) and EM [17] finds the maximum likelihood estimates. The read classification of Section 2.5 is based on the estimated proportions.

#### E-step

Given the current proportions, the E-step computes the posterior probability that read *i* was generated from leaf *l*,

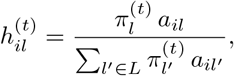

and sums these posteriors into an expected read count 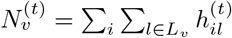 for every vertex *v*. These counts describe how reads divide at each vertex, so we parameterize the proportions the same way, by the conditional proportion *q*_*c*|*v*_ = *π*_*c*_*/π*_*v*_ of each child *c* of an internal vertex *v*, with ∑_*c∈ch*(*v*)_ *q*_*c*|*v*_ = 1. The proportion of a vertex is then the product of the conditionals along its root path (Figure 1d), the expected complete-data log-likelihood separates over internal vertices, and the M-step can maximize 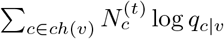 at each vertex *v* independently. See Supplementary Section S2 for the complete-data log-likelihood and the derivation of this decomposition.

#### Sparsity penalty

In many applications, few of the references are expected to be present in the sampled population. To impose this knowledge on the estimation, we penalize [42,41,9] each conditional proportion and maximize

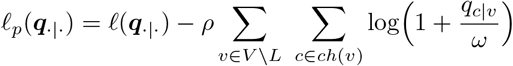

where *ρ* ≥ 0 sets the strength of the penalty and *ω >* 0 its scale. The M-step objective still separates over the internal vertices, into 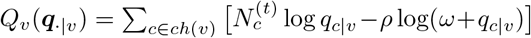 for each vertex *v*. See Supplementary Section S3 for the constrained penalized objective and the separation of the M-step.

#### M-step

For *ρ >* 0, we maximize a concave minorizer of *Q*_*v*_ that replaces the penalty with its tangent at the current estimate, which gives the update

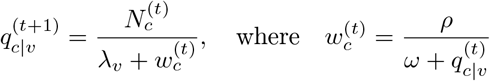

and the Lagrange multiplier *λ*_*v*_, found by Newton–Raphson, ensures the 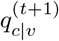 of the children of *v* sum to one. Without the penalty (*ρ* = 0), the update reduces to 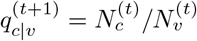, the share of *v*’s expected reads that fall under *c*. See Supplementary Section S3 for the minorizer, the derivation of the update, and the concavity of *Q*_*v*_.

#### Pruning

The M-step never returns an exact zero 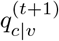 for a child with 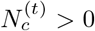, so unsupported vertices are removed explicitly: between the E-step and the M-step, every child with 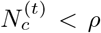, on which the penalized maximum places no mass, is deleted together with its subtree. With *ρ* = 0, no child is pruned. See Supplementary Section S3 for the derivation of this rule and for the case in which every child of a supported vertex is pruned.

### 2.4 Implementation

PREMISE restricts the sum in Eq. (2) to candidate alignments seeded by Super-Maximal Exact Matches of at least *κ* bases. It finds these seeds with the forward-backward algorithm of Li [25] on a run-length compressed bidirectional FM-index (BFM) [14,28,20,2] of all references. Each seed is extended over the full read and scored ungapped per Eq. (1), treating mismatches as base-call errors. A read whose best alignment likelihood is below *ϵ*_1_ is left unclassified, and leaf *l* is discarded as a source of read *i* if 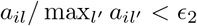.

EM starts from the taxonomy contracted to the references with read support (Figure 1c), with proportions initialized from each read’s maximum likelihood sources, and runs for a fixed number of iterations or until the penalized log-likelihood converges, updating the internal vertices in parallel. See Supplementary Section S4 and Supplementary Algorithm S1 for the full procedure and the default parameters.

### 2.5 Final Read Assignments

After EM, a final E-step gi ves each read’s posterior over leaves, 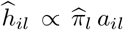, and summing over subtrees gives the probability 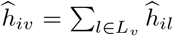 that read *i* originated below vertex *v*. Starting at the root, each read moves to its most probable child as long as that child’s posterior exceeds a confidence level *γ* ∈[0.5, 1), and is assigned to the vertex where it stops (Figure 1e). Reads that the references cannot distinguish below a given rank therefore remain at that rank instead of being forced to a leaf, and a read that stops at the root is reported as unclassified. See Supplementary Section S4 and Supplementary Algorithm S2 for the properties of this walk and its behavior without a taxonomy.

## 3 Experimental Setup

We conducted a comparison study that contrasts the performance of PREMISE with that of current state-of-the-art, general-purpose classification tools, namely Centrifuger v1.1.3 [19,49], Ganon v2.4.2 [44], Karp [45], KMCP v0.9.5 [48], Mora v1.0.0 [64], Salmon v2.5.1 [40] and Sylph v0.9.0 [47]. All computational experiments were conducted on a machine equipped with an Intel(R) Xeon(R) w7-2475X processor (40 cores, 80 threads), 128 GB of RAM, running Ubuntu 22.04.5 LTS (Linux kernel 5.15.0-173-generic). Methods were run one at a time with 40 threads, except for the KMCP search step, which used 4 threads as its documentation recommends. Each method was allowed one hour of wall-clock time per sample; KMCP exceeded it on all 30 synthetic influenza samples and on one real isolate. Methods that need a commercial solver or that report only taxa were not included. We compare the methods on dense samples, in which most references in the database are present, and on sparse samples, in which a few of many highly homologous references are present (see full sample compositions in Supplementary Table S1). PREMISE was run with the default seed length (*κ* = 22) and filters (*ϵ*_1_ = 0, *ϵ*_2_ = 10^*−*18^) in both regimes. On dense samples, it was given the NCBI taxonomy, run without the penalty (*ρ* = 0), and assigned reads at *γ* = 0.5. On sparse samples, it was run without a taxonomy and with the penalty (*ρ* = 150, *ω* = 10^*−*10^), and assigned reads at the default *γ* = 0.5. EM ran for a fixed 2,000 iterations on both.

### Dense samples

We used the first ten of the 100 short-read samples of the CAMI II strain-madness challenge [32] (samples 0–9, from the CAMI data portal [33]). Each contains 6.65 million read pairs (2 × 150 bp) simulated from 405 to 408 drawn from a pool of 408 genomes, many of them strains of the same species, with the true source of every read known. The 408 genomes (1.45 Gbp in 35,225 contigs) map to 26 NCBI taxa, 20 at the species rank and the others at the genus, family, or order. Every method was given the same reference of one record per genome, its contigs concatenated, and the methods that take a taxonomy (PREMISE, Centrifuger, and Ganon) the NCBI taxonomy, in which PREMISE attaches each genome as a leaf below its taxon (473 vertices). The true label of a read is its genome at the strain rank and each implied rank from kingdom to species (Supplementary Section S5.1).

### Sparse samples

The influenza reference comprises 5,268 Influenza A virus segments: all IAV sequences with complete assemblies in NCBI Virus [7] as of August 8, 2026, after removing old versions, sequences with more than 20% ambiguous nucleotides, exact duplicates, and contained sequences. We simulated three synthetic scenarios of ten samples each from database viruses with InSilicoSeq v2.0.1 [23] (500,000 MiSeq read pairs of 301 bp). The scenarios are isolates (Isolate-1 to Isolate-10), mixtures of three viruses of different subtypes (Subtype-1 to Subtype-10), and mixtures of three strains of one subtype (Strain-1 to Strain-10). Two real scenarios comprise four avian influenza isolates [15] and four defined mixtures of the PR8 and WSN33 H1N1 strains at ratios from 95/5 to 99.37/0.63 [30]. After primer and adapter trimming with cutadapt [29], the read pairs that BWA-MEM2 [56] aligns to a single known segment define the estimated ground truth. Because the closest absent references to the real segments are far less similar than co-occurring synthetic strains, we added two near-identical decoys of each of the 48 true segments of the real samples to the reference of every method. Details of the decoy generation procedure are provided in Supplementary Section S5.2.

### Evaluation metrics

At the read level, precision is the proportion of classified reads that are classified correctly, and coverage the proportion of input reads that are classified. Recall is the proportion of input reads classified correctly, and F1 the harmonic mean of precision and recall. For the real samples, whose ground truth is an estimate, we report coverage instead of recall. A reference is identified if its estimated abundance is positive; we count false positive (FP) and false negative (FN) references and compute the Sørensen similarity between the predicted and true reference sets. Abundance accuracy is the Ruzicka similarity between 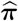 and ***π*** [58]; for Centrifuger, Ganon, and Mora, 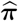 is computed from read assignment counts. On the dense samples, every metric is computed at each rank from kingdom to species, projecting truth and predictions up the taxonomy, and at the strain rank on genomes. A read classified above rank counts as unclassified at rank (Supplementary Section S5.3).

## 4 Results

We evaluate classification, profiling, runtime, and memory usage on the dense and then the sparse samples, and summarize index construction and parameter sensitivity.

### 4.1 CAMI II Comparative Analysis

All eight methods completed every sample. From kingdom to family, every method that classifies reads assigns nearly all of them correctly (precision ≥ 0.999 and coverage ≥ 0.998, except KMCP’s coverage of 0.988), whereas the abundance profiles of KMCP and Karp already deviate at the phylum (Ruzicka similarity 0.680 and 0.669). The methods separate at the genus and species. At the species, PREMISE has the highest precision (0.9986; 0.9967–0.9994) and coverage (98.72%), and hence the highest F1 score (0.992). It is followed by Ganon (0.980), Mora (0.971), KMCP (0.959), Salmon (0.956), and Centrifuger (0.879), whose coverage falls to 78.98% as it places reads with tied hits at their lowest common ancestor. PREMISE, Centrifuger, Ganon, Mora, and Salmon detect exactly the species present (Sørensen similarity 1.0; no FP or FN at any rank down to the species), while KMCP and Sylph miss 1.9 and 3.1 species per sample on average. Salmon estimates species abundances most accurately (Ruzicka similarity 0.978), followed by PREMISE (0.966) and Mora (0.955); Ganon (0.909), Sylph (0.881), Centrifuger (0.756), KMCP (0.660), and Karp (0.252) trail. The difference between PREMISE and Salmon lies in one pair of taxa: PREMISE assigns 2–5% of each sample’s abundance from genomes labeled only with the family Enterobacteriaceae to *E. coli* genomes, with which they share 17% of their sequence verbatim (17 Mbp).

At the strain rank, each read must be traced to the genome it was simulated from among up to 174 of the same species (Figure 2, right-most axes). Ganon, Mora, Salmon, and KMCP assign nearly every read to a genome, but only 32.6%, 18.1%, 15.9%, and 15.5% of reads to the correct one, so their precision equals their recall. PREMISE has the highest strain precision (0.804; 0.732–0.862) and F1 score (0.493, against 0.326 for Ganon), assigning 44.8% of the reads to a genome and the rest to their species. Centrifuger, which places reads with tied hits at their lowest common ancestor, assigns 15.0% of the reads to a genome at a precision of 0.639. PREMISE and Salmon estimate genome abundances far more accurately than the other methods (Ruzicka similarity 0.681 and 0.680, against at most 0.283 for Mora), with each ahead on five of the ten samples. Neither PREMISE nor Centrifuger nor Mora misses a genome present in a sample, and Salmon misses 0.8 per sample, whereas Ganon, Karp, KMCP, and Sylph miss on average 51, 80, 351, and 376 of the 405 to 408 present genomes (Supplementary Figure S1). Every method completed each CAMI II sample within about 20 minutes; PREMISE took 572 s on average (546–594 s), after Salmon (531 s) and before Karp (845 s), and needed 45.9 GB of memory, second to Mora (52.9 GB; Supplementary Figure S2).

**Fig. 2:**
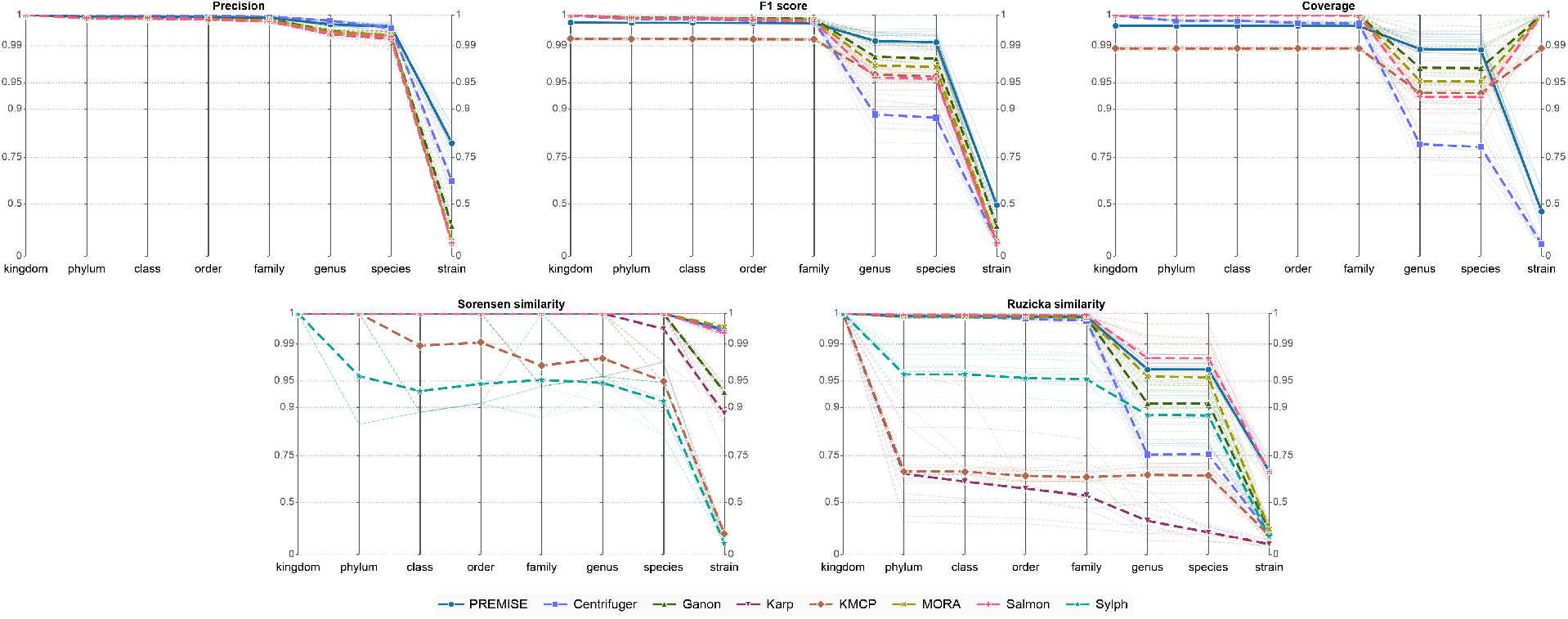
Classification and profiling accuracy across taxonomic ranks on CAMI II. Parallel coordinates of the precision, F1 score, coverage, Sørensen similarity, and Ruzicka similarity of each method at every rank from kingdom to species and, on the right-most axis, at the strain rank, whose labels are CAMI’s genomes. Each vertical axis is arctan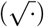-scaled from 0 (bottom) to 1 (top). Faint lines are the ten samples, bold lines with markers the means. Karp and Sylph do not classify reads, so they appear only in the Sørensen and Ruzicka panels. PREMISE assigns reads at *γ* = 0.5.

**Fig. 3:**
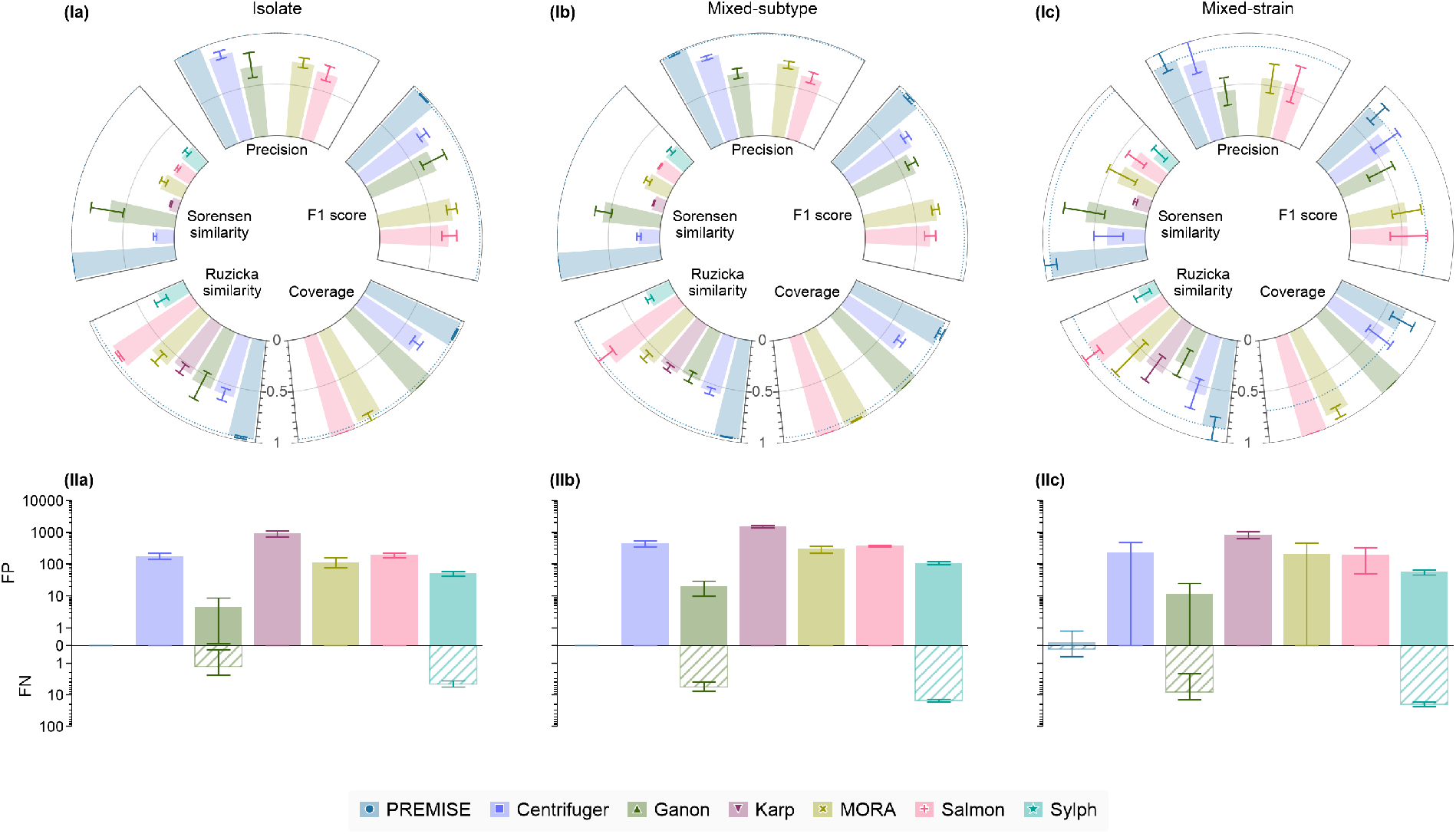
Inferential performance on synthetic data. Three columns show results for the synthetic viral isolate, mixed-subtype, and mixed-strain samples (ten samples each). The top radial plots (Ia–Ic) compare (clockwise from the top) the arcsine square root scaled precision, F1 score, coverage, Ruzicka similarity, and Sørensen similarity of the methods; the dotted arc marks the mean of PREMISE. The bottom butterfly plots (IIa–IIc) report the log-scaled number of false positive (FP, above the horizontal axis) and false negative (FN, below) reference identifications. Bars are means over the ten samples and whiskers one standard deviation. Karp and Sylph do not classify reads, so they have no precision, coverage, or F1 bars, and KMCP did not finish any synthetic sample within the one-hour limit (Section 3).

**Fig. 4:**
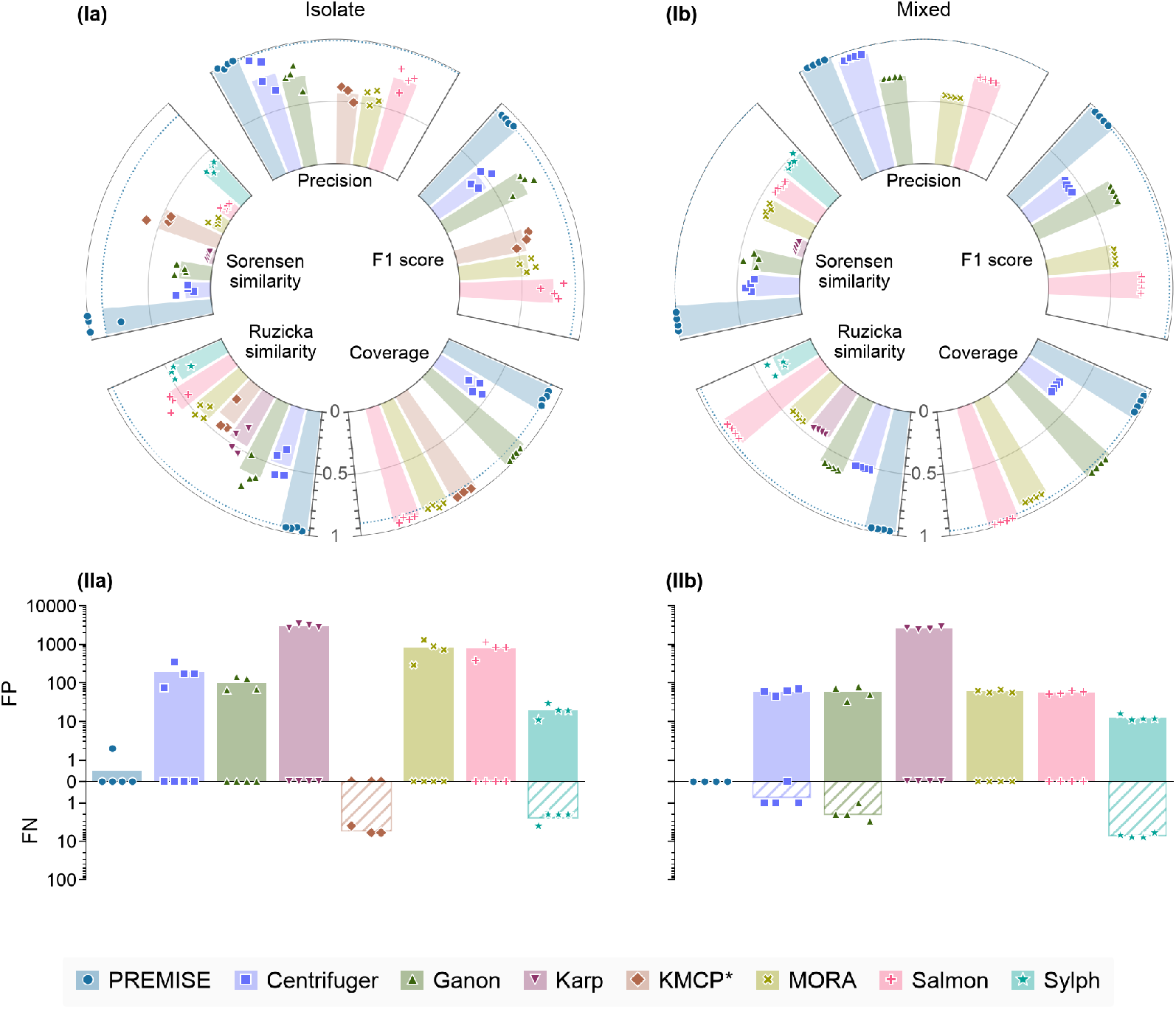
Inferential performance on real data. Two columns show results for the real viral isolate and mixed infection samples (four samples each). Panels are as in Figure 3, except that bars are means and each point is one sample. KMCP (asterisk) did not finish one isolate (SRR31013465) within the one-hour limit and returned an empty abundance profile for every mixed infection, so it is absent from (Ib) and (IIb).

### 4.2 Influenza Comparative Analysis

#### Synthetic samples

On the synthetic isolates, PREMISE recovers exactly the true references of all ten samples, with perfect precision (1.0), coverage of 99.44%–99.86%, and Ruzicka similarity ≥0.992. On the subtype mixtures, it recovers exactly the true references of all ten samples (Sørensen similarity 1.0), at a read precision of 0.997–1.000 and coverage of 98.33%–99.80%. On the strain mixtures, the hardest scenario, it keeps precision at 0.849–0.998 with no false references in nine samples and misses one reference each in Strain-1 and Strain-6, but leaves more reads unclassified (coverage 55.20%–99.26%, mean 77.77%). PREMISE has the highest readlevel precision on every synthetic sample except Strain-1, Strain-5, Strain-6, and Strain-7, where Centrifuger is more precise but classifies only 24.85%–86.30% of the reads. Restricted to the reads that Centrifuger classifies, PREMISE is the more precise of the two on nine of the ten strain mixtures (Barnard’s exact test [3], one-sided *p <* 10^*−*6^; *p* = 0.96 on Strain-3). The other methods trade precision for false references. Salmon assigns every read but reports 138 to 457 false references per sample, and Ganon’s precision falls to 0.163–0.824 on the strain mixtures. Centrifuger, Karp, and Mora report hundreds to thousands of false references (up to 1,784 for Karp), and Sylph misses three to six of the eight segments of every isolate. PREMISE and Salmon are the two most accurate abundance estimators: PREMISE has the highest Ruzicka similarity on nine, ten, and eight of the ten isolate, subtype-mixture, and strain-mixture samples, Salmon on the other two strain mixtures, and Ganon on Isolate-3.

#### Real samples

Across the eight real samples, PREMISE detects one near-identical decoy in one isolate and none in the other samples, whereas Ganon detects 7–12 per sample, Centrifuger 16–30, Salmon 24–33, Mora 26–33, and Karp 44–76. On the isolates, PREMISE maintains near-perfect read precision (1.00 on three isolates and 0.998 on the fourth) while assigning 96.7%–97.9% of reads. It recovers exactly the eight segments of three isolates, reports two false references on the fourth, and misses none. It also has the highest Ruzicka similarity of all methods on every isolate (mean 0.995). On the mixtures, it recovers the 16 true segments of both strains in every sample with no false positives or negatives, a read precision of 1.0, 99.5%– 99.6% of reads assigned, and a mean Ruzicka similarity of 0.997. Centrifuger keeps high precision (mean 0.89 on the isolates and 0.99 on the mixtures) but assigns only 21%–43% of reads and reports 45–340 false references per sample. Ganon assigns nearly all reads at a precision of 0.66–0.92, with 33–142 false references. Salmon assigns 99.9% of the mixture reads at a precision of 0.84–0.85, with 52–62 false references, and is the next best abundance estimator there (mean Ruzicka similarity 0.98). KMCP misses four to six of the eight segments of the isolates it completed and returns an empty profile for every mixture.

#### Resource utilization

On the influenza samples, PREMISE took 13.1–39.9 s and 0.73–2.53 GB per sample, 4–7× less memory than Mora in every scenario. Its mean runtime is below those of Salmon and Mora on the synthetic samples; on the real samples, Mora is faster on the isolates (13.2 vs. 16.4 s) and Salmon on the mixtures (10.9 vs. 30.8 s). Sylph completes in under a second, Centrifuger within 5 s, and Ganon and Karp within 36 s, all using under 1.8 GB, so PREMISE needs about an order of magnitude more time and memory than the sketching tools. KMCP did not finish any synthetic sample within the one-hour limit (Supplementary Figure S3).

### 4.3 Index Construction and Parameter Sensitivity

PREMISE builds its index in 1.57 s for the influenza reference (45.25 MB) and in 253 s for the 160 times larger CAMI II reference, whose index grows only 40-fold, to 1.83 GB (Supplementary Section S6). A one-at-a-time sweep of the six parameters of PREMISE on the real samples (Supplementary Section S8) leaves precision and Jaccard similarity unchanged for seed lengths *κ* ∈[20, 24], *ϵ*_1_ up to 10^*−*128^, *ϵ*_2_ from 10^*−*36^ to 10^*−*9^, and *ω* from 10^*−*30^ to 10^*−*6^. The penalty *ρ* trades false references against abundance accuracy; with 2,000 EM iterations, *ρ* = 150 recovers exactly the true segments of all mixtures and of three of four isolates.

## 5 Discussion

PREMISE accelerates full-read alignment with a run-length compressed bidirectional FM-index (BFM) enough to afford the likelihood calculations of a Phred-aware read model. It estimates the reference proportions by EM over a taxonomy, optionally penalized to encourage sparsity, thereby ignoring rare contaminants and artifacts, and assigns each read to the deepest taxon the data confidently support.

The two regimes we studied require different estimators. On the dense CAMI II samples, where nearly every reference is present, the maximum likelihood estimate (*ρ* = 0) detected exactly the species present at every rank, as did Centrifuger, Ganon, Mora, and Salmon. On the sparse influenza samples, where a few of many near-identical references are present, most other methods reported tens to thousands of false references, including dozens of the near-identical decoys on each real sample. KMCP avoided them only by also discarding true sources. The penalty (*ρ >* 0) kept PREMISE to four false references across all 38 samples while missing only two true sources. Judicious taxonomic placement helps in the dense regime. Rather than force a read onto one of several near-identical strains or onto their lowest common ancestor regardless of the evidence, as Centrifuger does, PREMISE stops each read at the deepest vertex supported by the posterior probability. It therefore classifies 98.7% of the reads at the CAMI II species rank, against 79.0% for Centrifuger, at a higher precision. The same rule sets PREMISE apart at the strain rank, where methods that force genome-level calls make incorrect assignments for over two-thirds of reads. PREMISE assigns fewer reads to a genome, but makes four in five correct assignments and recovers every genome present. Near-identical strains with disparate labels remain difficult to resolve. On CAMI II, PREMISE confuses read assignments between genomes sharing 17 Mbp identical sequence, some assigned to *E. coli* species and others resolved only to family, but Salmon apportions reads more accurately in nine of the ten samples.

Among methods that optimize probability models, PREMISE and Salmon estimate abundances most accurately, but Salmon, designed for RNA-seq quantification, has no sparsity mechanism and reports hundreds of false references on the synthetic samples and 24 to 33 decoys on each real one. Of the heuristic methods, Ganon over-calls references in sparse samples, Centrifuger maintains precision by leaving many reads at lowest common ancestors, and KMCP’s filters true sources with the false ones (Supplementary Section S7).

PREMISE advantages come with a main trade-off of low coverage: 45% of reads are left unclassified in the synthetic mixed-strain samples. These requirements make PREMISE better suited to the high resolution identification and monitoring of well-characterized pathogens and defined communities than to ultra-high-throughput, real-time screening of massive datasets. Future work will cut the memory needed for dense samples with equivalence classes of reads [24,5] and estimate the error and coverage properties from the data [8]. It will also be extended to detect reads from novel variants that violate the completeness assumption [45], apply PREMISE to real, complex metagenomes such as wastewater, and model indels with a pair hidden Markov model for noisier sequencing technologies.

## 6 Data availability

The source code for PREMISE is publicly available on GitHub at https://github.com/sriram98v/premise under a MIT license. The datasets used in the comparative study, including simulated viral sequences and empirical metagenomic data, are archived and available in the Zenodo repository at https://doi.org/10.5281/zenodo.19026876. The CAMI II strain-madness samples, source genomes, and taxonomy are available from the CAMI data portal [33].

## 8 Acknowledgments

This study was supported by an Iowa State University Research and Innovation Roundtable award and the Agricultural Research Service, United States Department of Agriculture (USDA-ARS project numbers 5030-32000-231-000D and 5030-32000-231-095-S); the National Institute of Allergy and Infectious Diseases, National Institutes of Health, Department of Health and Human Services (Contract No. 76N93021C00015); the SCINet project of the USDA-ARS (USDA-ARS project number 0500-00093-001-00-D); and the Iowa Agriculture and Home Economics Experiment Station, Ames, Iowa (project number IOW05806 supported by USDA/NIFA and State of Iowa funds). The funders had no role in study design, data collection and interpretation, or the decision to submit the work for publication. Mention of trade names or commercial products in this article is solely for the purpose of providing specific information and does not imply recommendation or endorsement by the USDA. USDA is an equal opportunity provider and employer.

