## Supplement for "PREMISE: A Quality-Aware Probabilistic Framework for High-Resolution Source Assignment in Metagenomic Sequencing"

### S1 Related Work

Profiling in the presence of complete databases was first attempted in RNA-seq, where near complete transcript annotations allow modeling of reads as sequencing variation around known transcripts. RSEM [22] uses the Expectation Maximization (EM) algorithm to obtain Maximum Likelihood Estimates (MLEs) of transcript abundances under a model for sequenced transcript fragments, assuming the top scoring alignments found by an independent aligner identify the only possible sources of each fragment. Efficiency gains result from grouping read sequences into equivalence classes of possible source transcripts and replacing reads with class counts. Kallisto [4] pseudoaligns reads to equivalence classes via $k$-mer lookups and ignores all other differences between reads in a class, while Salmon [34,30] restores model detail by modeling some read biases. Metakallisto [38] and Salmon option `--meta` confirm such quantification can work for metagenomes.

Metagenomics tools are, however, mostly designed to solve taxonomic classification and profiling against incomplete databases [28] of reference sequences registered to leaf nodes of a hierarchical taxonomy. In this context, probability models must explain both evolutionary variation around the known references and measurement error by sequencing. Sigma [1] was the first metagenomic tool to pose a probability model. It uses nonlinear programming to find MLE abundances assuming independent and identically distributed (iid) nucleotide model with fixed substitution rate (default 5%) and no penalty for insertions and deletions (indels). Karp [37] implicitly assumes a complete database and uses accelerated EM to find abundance MLEs with sequencing errors explained by Phred error probabilities [11] and no indel penalties, but it identifies candidate alignments by initial pseudoalignment [4] adjusted to tolerate sequencing errors. Mora [50] takes the top alignments and their scores from a sequence aligner, finds the MLE abundance estimates via EM supplemented with AGAMEMNON [41] iterations to drop poorly supported references, and reassigns reads to references via post-hoc application of the weapon-target assignment problem. Sylph [39] splits read variation into evolutionary substitutions and sequencing errors, both assumed iid, but operates on $k$-mer sketches for computational efficiency. It uses a zero-inflated Poisson model over $k$-mer multiplicities to estimate containment Average Nucleotide Identity (ANI) [14] and effective coverage, reassigning shared $k$-mers to profile only sampled species. All statistical methods compromise, usually in the probability model, to account for incomplete databases or retain computational efficiency.

Most metagenomic methods do not use an explicit probability model, but achieve validity through judicious thresholds and filters [49]. Such metagenomic methods use variable amounts of linkage information in the read to improve resolution. From methods that treat reads as bags of $k$-mers to methods that align whole reads, there is general trade-off of speed for resolution.

The $k$-mer methods tend to be fast. Kraken [46] assigns read $k$-mers to the Lowest Common Ancestor (LCA) of containing taxa and reads to the LCA hosting more of its $k$-mers of any leaf-to-root path. Kraken 2 [45] inserts a subset of $k$-mers into a compact hash table to reduce time and space complexity. Both versions of Kraken filter taxa by number of assigned reads, but KrakenUniq [5] filters by an estimated count of unique $k$-mers witnessed in the reads. CLARK [33] indexes only $k$-mers unique to each reference and assigns reads by most matching $k$-mers; CLARK-S [32] modifies CLARK to use spaced seeds. Bracken [25] redistributes Kraken counts up and down the taxonomy through one EM iteration, initialized with Kraken taxon abundances and assuming uniform read coverage with no read errors. While Ganon [35] assigned reads to the reference with the most matched $k$-mers, Ganon2 [36] models reads as partially observed multinomial draws, arising from any reference with sufficient matched $k$-mers, then uses hard EM to estimate multinomial probabilities. Both achieve speed by storing $k$-mers in interleaved Bloom filters of references partitioned within taxonomic levels. KMCP [40] looks up $k$-mers in a modified compact bit-sliced signature index with flexible genome chunking for large databases and fine control over the sensitivity-specificity-memory trade-off. It also filters candidate references, requiring sufficient uniquely and well-aligned reads, uniform read coverage, and that no other reference can explain the assigned reads [21], finally redistributing reads of discarded references.

Methods utilizing partial or full read linkage offer varying balances of speed and precision. MEGAN [16] recruits BLASTN [7] to locally align reads to references and assigns reads to the LCA of references with accepted alignments [15], redistributing multiply aligned reads down the hierarchy in proportion to abundances estimated from uniquely aligned reads. Centrifuge [17] discards reference fragments $\geq$ 99% similar to other genomes before loading the rest into a substantially smaller FM index. Then, starting with the 5' ends of the read and its reverse complement, it extends $k$-mer matches, restarting with the $k$-mer after each failed lookup, empirically scores hits based on match lengths, and assigns reads to $\leq 5$ equally highest scoring references by greedily substituting LCAs. Metalign [19] filters candidate sources from a large reference database by computing the containment MinHash similarity metric [18] to ensure a minimum proportion (default: 0.01) of a retained reference $k$-mers are covered in the reads. Keeping one strain per species, it then uses Minimap2 [24] to align reads to selected species representative sequences. Finally, retaining only species with at least one uniquely mapped read, it estimates abundances in one iteration of EM, initialized with estimates based on uniquely mapped reads and assuming equal likelihoods for all found alignments consuming 95% or more of the read. CAMMiQ [51] indexes shortest variable-length substrings unique to one or two reference genomes into a compact trie structure. It formulates a mixed integer linear program assuming uniform coverage, iid errors, and user-guided constraints on abundance to identify the sampled references and their abundances. Like KMCP, CAMMiQ heavily filters references, requiring at least one confidently mapped read, at least 10% uniquely mapped reads, reads covering a substantial proportion of strain-unique subsequences, and insubstantial difference between average strain-specific subsequence coverage and overall estimated strain coverage. PanTax [48] uses a read's optimal alignment to a pangenome graph capturing intraspecies genetic variation. Strain abundances are estimated by minimizing absolute difference between observed and implied node abundances using two passes of integer linear programming. Candidate strains are filtered by read coverage of strain-unique nodes to require minimum coverage before the first pass and minimal disagreement with first-pass estimated strain coverage before the second pass.

Several benchmarks of metagenomic tools offer confusing recommendations, but reveal consistent flaws: false positives remain a serious problem, and there is a dramatic drop-off in performance at the resolution of species or higher [47,8,28,42,29]. Among them, the CAMI challenges [28,29] have become the reference standard, scoring classification and profiling at every taxonomic rank against the known origin of each simulated read, and their strain-madness samples remain among the hardest for profilers. Therefore, there remains a need for high resolution taxonomic classification and profiling, particularly for the enriched metagenomic samples used in diagnostics for well-characterized viral pathogens.

### S2 Complete-Data Log-Likelihood over the Taxonomy

This section derives the complete-data log-likelihood of the read data $\mathcal{R} = \{\boldsymbol{r}_i\}_{i=1}^n$ and its decomposition over the taxonomy, using the notation of Section 2.1. It utilizes the read likelihood conditional on source and alignment $a_{ilz} = \Pr(\boldsymbol{R}_i = \boldsymbol{r}_i \mid S_i = l, Z_i = z)$, the read likelihood conditional on source $a_{il} = \Pr(\boldsymbol{R}_i = \boldsymbol{r}_i \mid S_i = l)$, and the read $i$ alignment start probability $\eta_{ilz} = Pr(Z_i = z \mid S_i = l)$ given source $l$ proposed in Section 2.2 of the main text.

*Data log-likelihood.* Assuming the $n$ reads are generated independently, the observed-data log-likelihood is

$$\ell(\pi) = \sum_{i=1}^{n} \log \sum_{l \in L} \pi_l \, a_{il} = \sum_{i=1}^{n} \log \Big[ \sum_{l \in L} \pi_l \sum_{z=1}^{|\boldsymbol{D}_l| - |\boldsymbol{r}_i| + 1} \eta_{ilz} \, a_{ilz} \Big].$$

Because we assume uniform read coverage within source $\boldsymbol{D}_l$, the unknown alignment start position $Z_i$ can be marginalized out in read likelihood $a_{il}$, but the sum over leaves makes $\ell$ difficult to maximize directly. So, we treat the source leaf $S_i$ of each read as missing data and formulate the complete-data log-likelihood as

$$\ell_c(\pi) = \sum_{i=1}^{n} \sum_{l \in L} \mathbb{1}(S_i = l) \big[ \log \pi_l + \log a_{il} \big].$$

*Decomposition over the taxonomy.* Given the proportions $\boldsymbol{\pi}^{(t)}$ at EM iteration $t$, the E-step computes the posterior probability $h_{il}^{(t)} := \mathbb{P}(S_i = l \mid \boldsymbol{r}_i; \boldsymbol{\pi}^{(t)}) = \pi_l^{(t)} a_{il} \big/ \sum_{l' \in L} \pi_{l'}^{(t)} a_{il'}$ that read $i$ was generated from leaf $l$, and the expected read count $N_v^{(t)} = \sum_{i=1}^n \sum_{l \in L_v} h_{il}^{(t)}$ of every vertex $v$. The counts respect the tree structure: $N_r^{(t)} = n$, and $N_v^{(t)} = \sum_{c \in ch(v)} N_c^{(t)}$ for every internal vertex $v$. For each internal vertex $v$ and child $c \in ch(v)$, the conditional proportion $q_{c|v} = \pi_c / \pi_v$ satisfies $q_{c|v} \geq 0$ and $\sum_{c \in ch(v)} q_{c|v} = 1$, and the proportion of any vertex is the product of the conditionals along its root path, $\pi_v = \prod_{(p,c) \in E(rTv)} q_{c|p}$, where $E(rTv)$ is the edge set of the path. Taking the expectation of $\ell_c$ and writing $\log \pi_l$ as the sum of the log-conditionals along $rTl$, each edge $(p, c)$ of $T$ collects the expected counts of every leaf in $L_c$:

$$\begin{aligned}
\sum_{i=1}^{n} \sum_{l \in L} h_{il}^{(t)} \big[\log \pi_l + \log a_{il}\big] &= \sum_{l \in L} N_l^{(t)} \log \pi_l + C \\
&= \sum_{l \in L} N_l^{(t)} \sum_{(p,c) \in E(rTl)} \log q_{c|p} + C \\
&= \sum_{(p,c) \in E(T)} \log q_{c|p} \sum_{l \in L_c} N_l^{(t)} + C \\
&= \sum_{v \in V \setminus L} \sum_{c \in ch(v)} N_c^{(t)} \log q_{c|v} + C,
\end{aligned}$$

where $E(T)$ is the directed (root to leaf) edge set of $T$, and $C = \sum_{i=1}^n \sum_{l \in L} h_{il}^{(t)} \log a_{il}$ does not depend on $\boldsymbol{\pi}$.

### S3 Penalized Objective

The M-step maximizes a minorizer of a penalized objective and prunes unsupported children. This section states the objective and shows why both steps are needed.

*Penalized objective.* The penalized log-likelihood

$$\ell_p(\boldsymbol{q}_{\cdot|\cdot}) = \ell(\boldsymbol{q}_{\cdot|\cdot}) - \rho \sum_{v \in V \setminus L} \sum_{c \in ch(v)} \log\Big(1 + \frac{q_{c|v}}{\omega}\Big)$$

is maximized subject to $q_{c|v} \geq 0$ and $\sum_{c \in ch(v)} q_{c|v} = 1$ for every internal vertex $v$, where $\ell(\boldsymbol{q}_{\cdot|\cdot})$ is the log-likelihood with each $\pi_l$ reparameterized as a product of conditionals $q_{c|v}$, $\rho \geq 0$ sets the strength of the penalty, and $\omega > 0$ sets its scale. Adding the penalty to the expected complete-data log-likelihood (Section S2), the function maximized in the M-step is, up to a constant,

$$Q(\boldsymbol{q}_{\cdot|\cdot}) = \sum_{v \in V \setminus L} Q_v(\boldsymbol{q}_{\cdot|v}), \qquad Q_v(\boldsymbol{q}_{\cdot|v}) = \sum_{c \in ch(v)} \Big[N_c^{(t)} \log q_{c|v} - \rho \log\big(\omega + q_{c|v}\big)\Big],$$

where the constant $\rho \log \omega$ per child is dropped from $Q_v$. Each $Q_v$ depends only on the conditional proportions of the children of $v$ and is subject only to the sum-to-one constraint of $v$, so the M-step separates into one independent problem per internal vertex, and the vertices interact only through the E-step.

*Minorizer.* For $\rho > 0$, $Q_v$ need not be concave (see the concavity analysis below), so instead of maximizing $Q_v$ directly, we maximize a concave lower bound $G_v$ that replaces the convex penalty term with its tangent at the current estimate, a minorizer built from a supporting hyperplane:

$$G_v(q_{\cdot|v}; q_{\cdot|v}^{(t)}) = \sum_{c \in ch(v)} \Big[N_c^{(t)} \log q_{c|v} - w_c^{(t)} q_{c|v}\Big], \qquad w_c^{(t)} = \frac{\rho}{\omega + q_{c|v}^{(t)}}.$$

The function $\log(\omega + q)$ is concave, so its tangent lies above it; the tangent to $-\rho \log(\omega + q)$ therefore lies below it, and $G_v \leq Q_v$, with equality at $\boldsymbol{q}_{\cdot|v}^{(t)}$. Introducing a Lagrange multiplier $\lambda_v$ for the constraint $\sum_c q_{c|v} = 1$ and setting the derivative of the Lagrangian to zero gives the update

$$q_{c|v}^{(t+1)} = \frac{N_c^{(t)}}{\lambda_v + w_c^{(t)}}, \qquad \text{where } \lambda_v \text{ solves } \sum_{c \in ch(v)} \frac{N_c^{(t)}}{\lambda_v + w_c^{(t)}} = 1,$$

and PREMISE finds $\lambda_v$ by Newton–Raphson (Algorithm S1). Without the penalty ($\rho = 0$), $w_c^{(t)} = 0$, $G_v = Q_v$, and the update reduces to $q_{c|v}^{(t+1)} = N_c^{(t)}/N_v^{(t)}$.

*Concavity.* We now determine when $Q_v$ is concave, which decides whether it could be maximized directly.

**Theorem 1.** *$f$ is concave on $(0,1]$ $\iff \rho \le N(1+\omega)^2$.*

*Proof.* The Hessian of $Q_v$ is diagonal, because $Q_v$ is a sum of terms that each depend on a single child; $Q_v$ is therefore concave if and only if the term of every child is concave, and it suffices to study the term of an arbitrary child $c$ of $v$,

$$f_c(q) = N_c^{(t)} \log q - \rho \log(\omega + q), \qquad q \in (0,1],$$

whose derivatives are

$$f_c'(q) = \frac{N_c^{(t)}}{q} - \frac{\rho}{\omega + q}, \qquad f_c''(q) = \frac{\rho}{(\omega+q)^2} - \frac{N_c^{(t)}}{q^2}.$$

The term $f_c$ is concave where

$$f_c''(q) < 0 \iff \frac{\rho}{(\omega+q)^2} < \frac{N_c^{(t)}}{q^2} \iff q\left(\sqrt{\rho} - \sqrt{N_c^{(t)}}\right) < \omega\sqrt{N_c^{(t)}}.$$

Moving forward, we simply write $N$ for $N_c^{(t)}$.

For $q > 0$, both sides below are non-negative, so taking square roots preserves the inequality:

$$f''(q) \le 0 \iff \rho\, q^2 \le N(\omega+q)^2 \iff \sqrt{\rho}\, q \le \sqrt{N}\,(\omega+q) \iff q\big(\sqrt{\rho} - \sqrt{N}\big) \le \omega\sqrt{N}.$$

Write $d := \sqrt{\rho} - \sqrt{N}$, so that

$$f''(q) \le 0 \iff q\, d \le \omega\sqrt{N}. \tag{S1}$$

Also, since $\rho, N \ge 0$,

$$\rho \le N(1+\omega)^2 \iff \sqrt{\rho} \le (1+\omega)\sqrt{N} \iff d \le \omega\sqrt{N}. \tag{S2}$$

**Case 1:** $\Longleftarrow$

Assume $\rho \le N(1+\omega)^2$. By Eq. (S2), $d \le \omega\sqrt{N}$. Take any $q \in (0,1]$. There are two cases.

- $d \le 0$ (that is, $\rho \le N$). Then $q\, d \le 0 < \omega\sqrt{N}$, because $q > 0$, $\omega > 0$ and $N > 0$.
- $d > 0$ (that is, $N < \rho \le N(1+\omega)^2$). Then $q\, d \le 1 \cdot d \le \omega\sqrt{N}$, using $q \le 1$ and then $(**)$.

In both cases $q\, d \le \omega\sqrt{N}$, so $f''(q) \le 0$ by $(*)$. This holds for every $q \in (0,1]$, so $f$ is concave on $(0,1]$.

**Case 2:** $\Longrightarrow$

Prove the contrapositive: assume $\rho > N(1+\omega)^2$ and show that $f$ is not concave. By Eq. (S2), $d > \omega\sqrt{N} > 0$. At $q = 1$:

$$1 \cdot d > \omega\sqrt{N} \overset{(*)}{\Longrightarrow} f''(1) > 0.$$

$f''$ is continuous on $(0,1]$, so there is a $\delta > 0$ with $f'' > 0$ on $(1-\delta, 1]$. On that subinterval $f$ is strictly convex, so it cannot be concave on $(0,1]$. This contradicts the hypothesis.

More precisely, $f$ is concave on $(0, q^*]$ and strictly convex on $(q^*, 1]$, where $q^* = \omega\sqrt{N}/d < 1$.

□

*Pruning.* The M-step never returns an exact zero $q_{c|v}^{(t+1)}$ for a child with $N_c^{(t)} > 0$, so unsupported vertices must be removed explicitly. The pruning criterion follows from the behavior of the term of each child in $Q_v$ for small $\omega$: with $\log(1 + q/\omega) = \log(\omega + q) - \log\omega$, the term satisfies

$$N_c^{(t)} \log q_{c|v} - \rho \log\Big(\omega + q_{c|v}\Big) \longrightarrow \big(N_c^{(t)} - \rho\big) \log q_{c|v} \qquad \text{as } \omega \to 0.$$

When $N_c^{(t)} < \rho$, this limit increases as $q_{c|v} \to 0$, and the stationarity condition $(N_c^{(t)} - \rho)/q_{c|v} = \lambda_v$ has no positive solution, so the penalized maximum places no mass on $c$. Pruning is therefore performed between the E-step and the M-step: every child with $N_c^{(t)} < \rho$ is deleted together with its subtree, and the M-step is solved over the remaining children. Counts aggregate up the tree, so $N_c^{(t)} < \rho$ implies that every vertex in the subtree of $c$ also falls below $\rho$, and deleting the whole subtree never removes a vertex that would have passed the test on its own. With $\rho = 0$, no child is pruned.

The tree introduces one case without a flat counterpart: every child of an internal vertex $v$ may fall below $\rho$ while $N_v^{(t)} \geq \rho$, which happens when the reads support $v$ but none of its descendants individually. Deleting all children of $v$ would leave the mass of $v$ without a leaf to hold it, so we instead move the reads of the children up to $v$ and omit the subtrees of all children of $v$ from the following EM iterations.

### S4 Implementation

The processing of a sample has three stages: computing the read likelihoods, preprocessing the taxonomy, and iterating EM. They are described below and formalized in Algorithm S1; the final read assignment is formalized in Algorithm S2.

*Computing read likelihoods.* In principle, the read–alignment likelihood $a_{ilz}$ can be evaluated at every start position of every reference, but most of these alignments carry numerous mismatches and contribute negligibly to read likelihood $a_{il}$. PREMISE therefore restricts $a_{il}$ to candidate alignments along Supermaximal Exact Matches (SMEMs), treating $a_{ilz}$ as zero elsewhere. It finds the SMEMs in a Bidirectional Run-Length FM-index (B-RLFM) [12,2,10] of all reference sequences $\mathcal{D}$, in which a reference assembled from several contigs, such as a draft genome, can be given as one record with its contigs concatenated (Section S5.1). Alignments are seeded with all SMEMs of at least $\kappa$ bases (default: $\kappa = 22$), which the B-RLFM enumerates with the forward-backward algorithm of Li [23]. Each seed is extended in both directions until it spans the entire read; seeds on the same diagonal are collapsed into one candidate, and seeds on distinct diagonals remain separate candidates. Each candidate $(l, z)$ is scored ungapped over the full read against reference $l$ at offset $z$, and its likelihood $a_{ilz}$ is computed by Eq. (1), treating every mismatch as a base-call error. An ambiguous base in the read or the reference contributes the average of $a_{ilz}$ over the nucleotides it stands for; an N contributes probability 1/4 regardless of quality. The candidates of each read are summed per leaf according to Eq. (2) to give $a_{il}$.

With large reference databases, a read can have seeds in homologous regions shared by many references, which results in numerous candidate alignments and a heavy computational burden; PREMISE reduces the time and memory required by eliminating unlikely sources of each read. It declares read $i$ unclassified if the likelihood of its best alignment, $\max_{l,z} a_{ilz}$, is below $\epsilon_1$ (default: $\epsilon_1 = 0$, i.e., not used). It then discards leaf $l$ as a possible source of read $i$, setting $a_{il} = 0$, if $a_{il} / \max_{l' \in L} a_{il'} < \epsilon_2$ (default: $\epsilon_2 = 10^{-18}$), and stores the surviving $a_{il}$ in a sparse $n \times m$ matrix.

*Taxonomy preprocessing.* Every reference sequence $\boldsymbol{D}_l$ is attached as a leaf child of the most specific vertex to which it is resolved; references without a taxon, and all references when no taxonomy is given, are attached to the root. Taxonomic leaves that receive references thereby become internal vertices, so the leaf set of $T$ is exactly the reference set, whose members are individual genomes, strains, or segments. The leaf proportions are initialized from the maximum likelihood sources of each read, splitting a read evenly among tied sources, and the taxonomy is contracted to the references with a positive initial proportion, together with all vertices on their root paths. Finally, each $q_{c|v}$ is computed in a bottom-up traversal of the tree.

*EM iterations.* EM iterates for $t = 0, 1, 2, \ldots$ until a stopping rule is met. PREMISE either runs a fixed budget of $t_{\max}$ iterations or, after at least 50 iterations, stops once the relative change in the penalized log-likelihood $\mathcal{L}_P$ (Algorithm S1) between consecutive iterations falls below a threshold $\delta$ (default: $\delta = 10^{-6}$); an alternative rule monitors the change in the leaf proportions instead. We use the fixed budget of $t_{\max} = 2{,}000$ iterations for both dense and sparse samples (Section S7) and in the ablation study (Section S8). Each iteration consists of three steps.

E-step: The posteriors $h_{il}$ (Section S2) of each read $i$ are computed over the leaves $l$ with $a_{il} > 0$, and the expected counts are accumulated in a bottom-up traversal of the taxonomy, with $N_l^{(t)} = \sum_{i=1}^{n} h_{il}$ for every leaf $l$ and $N_v^{(t)} = \sum_{c \in ch(v)} N_c^{(t)}$ for every internal vertex $v$.

Prune: The taxonomy is traversed top-down, and at each internal vertex $v$, the children that exceed the penalty threshold form $C_v = \{c \in ch(v) : N_c \geq \rho\}$. If $C_v = \varnothing$, the subtree at $v$ is collapsed into a single leaf; otherwise, every child of $v$ outside $C_v$ is deleted. A pruned vertex is always removed by its parent before it is visited, so the collapse case is reached only by vertices with $N_v \geq \rho$. If the root is pruned to a single leaf, $\rho$ is too large for the sample.

M-step: Each internal vertex $v$ is updated independently, in parallel, using its surviving children $C_v$. If $v$ has a single child $c$, then $q_{c|v}^{(t+1)} = 1$; otherwise, the weights $w_c$ are computed from the current $q_{c|v}^{(t)}$, the Lagrange multiplier $\lambda_v$ is found, and $q_{c|v}^{(t+1)}$ is updated (Section S3).

**Algorithm S1** Hierarchical EM with optional penalty

**Require:** Reads with quality scores; reference database $\mathcal{D}$ and taxonomy $T$; minimum SMEM length $\kappa$; filtering thresholds $\epsilon_1 \geq 0$, $\epsilon_2 \geq 0$; penalty $\rho \geq 0$ (0 disables it); scale $\omega > 0$; iteration budget $t_{\max}$; optional stopping threshold $\delta > 0$

**Ensure:** Pruned taxonomy $T$; conditional proportions $\widehat{q}$; vertex proportions $\widehat{\pi}_v$

*Read alignment*

**for all** reads $i$ **do**
  Seed with all SMEMs of length $\geq \kappa$; extend each seed over the full read; collapse seeds sharing a diagonal
  Score each candidate $(l, z)$ ungapped over the full read to get $a_{ilz}$
  **if** $\max_{l,z} a_{ilz} < \epsilon_1$ **then** mark read $i$ unclassified and **continue**
  $a_{il} \leftarrow \sum_z \eta_{ilz}\, a_{ilz}$ over the candidates of each leaf
  $a_{il} \leftarrow 0$ for every $l$ with $a_{il} / \max_{l'} a_{il'} < \epsilon_2$; store the rest in a sparse matrix
**end for**

*Taxonomy preprocessing*

Attach each reference $\boldsymbol{D}_l$ as a leaf child of the most specific vertex it is resolved to
$M_i \leftarrow \arg\max_{l \in L} a_{il}$ for every read $i$; $\pi_l \leftarrow \frac{1}{n} \sum_{i=1}^{n} \mathbf{1}[l \in M_i] / |M_i|$ for every leaf $l$
Contract $T$ to the references with $\pi_l > 0$, together with their ancestors
$\pi_v \leftarrow \sum_{c \in ch(v)} \pi_c$ bottom-up; $q_{c|v} \leftarrow \pi_c / \pi_v$ for every edge $(v, c)$

*EM iterations*

**for** $t = 0, 1, \ldots, t_{\max} - 1$ **do**
  $h_{il} \leftarrow \pi_l\, a_{il} \big/ \sum_{l' \in L} \pi_{l'}\, a_{il'}$ for all $i$, and $l$ with $a_{il} > 0$ ▷ E-step
  $N_l \leftarrow \sum_{i=1}^{n} h_{il}$ for every leaf; $N_v \leftarrow \sum_{c \in ch(v)} N_c$ bottom-up
  $\mathcal{L}_P^{(t)} \leftarrow \sum_i \log \sum_l \pi_l\, a_{il} - \rho \sum_{v \in V \setminus L} \sum_{c \in ch(v)} \log\left(1 + q_{c|v}/\omega\right)$
  **if** $\delta$ is given **and** $t \geq 50$ **and** $\left|\mathcal{L}_P^{(t)} - \mathcal{L}_P^{(t-1)}\right| < \delta \max\left(1, \left|\mathcal{L}_P^{(t)}\right|\right)$ **then break**
  **for all** internal $v$, top-down, skipping deleted vertices **do** ▷ Prune
    $C \leftarrow \{c \in ch(v) : N_c \geq \rho\}$
    **if** $C = \emptyset$ **then**
      Collapse the subtree at $v$ into a single leaf
    **else**
      Delete every child $c \notin C$ together with its subtree
    **end if**
  **end for**
  **for all** internal $v$, in parallel, with children $C = ch(v)$ **do** ▷ M-step
    **if** $|C| = 1$ **then** $q_{c|v} \leftarrow 1$ and **continue**
    $w_c \leftarrow \rho / \left(\omega + q_{c|v}\right)$ for $c \in C$
    $k \leftarrow \arg\min_{c \in C} w_c$; $\lambda_v \leftarrow N_k - w_k$ ▷ Newton–Raphson
    **repeat**
      $g \leftarrow \sum_{c \in C} N_c / (\lambda_v + w_c)$; $g' \leftarrow -\sum_{c \in C} N_c / (\lambda_v + w_c)^2$
      $\lambda_v \leftarrow \lambda_v - (g - 1)/g'$
    **until** $\lambda_v$ converges
    $q_{c|v} \leftarrow N_c / (\lambda_v + w_c)$ for $c \in C$
  **end for**
  $\pi_r \leftarrow 1$; $\pi_c \leftarrow \pi_v\, q_{c|v}$ for every edge $(v, c)$, top-down
**end for**
**return** $T$, $\widehat{q}$, $\{\widehat{\pi}_v\}$

*Read assignment.* After EM, a final E-step gives the posterior $\widehat{h}_{iv} = \sum_{l \in L_v} \widehat{h}_{il}$ that read $i$ originated below vertex $v$, and each read walks down from the root while the posterior of its most probable child exceeds the confidence level $\gamma$; Algorithm S2 formalizes this walk. The posterior $\widehat{h}_{iv}$ can only decrease along any root-to-leaf path, with $\widehat{h}_{ir} = 1$, so the walk stops at the deepest vertex that the data confidently support. The confidence level $\gamma \in [0.5, 1)$ defaults to $\gamma = 0.6$. The posterior probabilities of the children of a vertex sum to at most one, so with $\gamma \geq 0.5$, at most one child qualifies, and the walk never has to break a tie. Without a taxonomy, every leaf is a child of the root, so a read is classified to its most probable reference if the posterior of that reference exceeds $\gamma$, and it is otherwise unclassified.

**Algorithm S2** Read assignment

**Require:** Pruned taxonomy $T$ and conditional proportions $\widehat{q}$ from Algorithm S1; read–leaf likelihoods $a_{il}$; confidence level $\gamma \in [0.5, 1)$
**Ensure:** An assigned vertex $v_i$ for every read $i$
$\widehat{\pi}_r \leftarrow 1$; $\widehat{\pi}_c \leftarrow \widehat{\pi}_v\, \widehat{q}_{c|v}$ for every edge $(v, c)$, top-down
**for all** reads $i$, in parallel **do**
    $\widehat{h}_{il} \leftarrow \widehat{\pi}_l\, a_{il} \big/ \sum_{l' \in L} \widehat{\pi}_{l'}\, a_{il'}$ for every leaf $l$ with $a_{il} > 0$ ▷ E-step
    $\widehat{h}_{iv} \leftarrow \sum_{c \in ch(v)} \widehat{h}_{ic}$ bottom-up over their ancestors; 0 elsewhere
    $v \leftarrow r$ ▷ $\widehat{h}_{ir} = 1$
    **while** $v \notin L$ **and** $\max_{c \in ch(v)} \widehat{h}_{ic} > \gamma$ **do**
        $v \leftarrow \arg\max_{c \in ch(v)} \widehat{h}_{ic}$
    **end while**
    $v_i \leftarrow v$ ▷ $v_i = r$: unclassified
**end for**
**return** $\{v_i\}_{i=1}^{n}$

### S5 Datasets and Preprocessing

This section details the benchmark samples summarized in Table S1: the dense CAMI II samples (Section S5.1), the sparse influenza samples and the decoy references added to their reference database (Section S5.2), and the evaluation metrics (Section S5.3).

#### S5.1 Dense Samples

In a dense sample, most references in the database are present, so the reads must be divided among near-identical strains that all contribute to the sample. The strain-madness samples of the CAMI II challenge [28] are the gold-standard example: hundreds of genomes from a few taxa, many of them strains of the same species, with the true source of every read known. Several of their genomes are resolved only to a genus, family, or order, so the samples also test how PREMISE handles references attached above the species rank.

*CAMI II strain-madness samples.* The challenge provides 100 short-read samples simulated from a common pool of 408 genomes; we used the first ten (samples 0–9), downloaded from the CAMI data portal [29]. Each sample contains 6.65 million read pairs ($2 \times 150$ bp) drawn from 405 to 408 of the 408 genomes. The interleaved read files were split into mate files, keeping the read identifiers that link each pair to the read-to-genome mapping of CAMI.

*Reference genomes.* The reference set comprises the 408 source genomes distributed with the challenge (1.45 Gbp in 35,225 contigs). We represented each genome as a single reference, with its contigs concatenated directly in file order, because CAMI simulates reads at the same depth on every contig of a genome. Of the contigs, 20% (7,018) occur verbatim in more than one genome; for example, the genomes labeled only with the family Enterobacteriaceae share 17% of their sequence (17 Mbp) verbatim with *E. coli* genomes. Every method was given the same reference set of one record per genome.

*Taxonomy.* CAMI assigns each genome an NCBI taxon identifier; the 408 genomes map to 26 taxa, 20 at the species rank and the others at the genus (3), family (2), and order (1) ranks. The taxa are highly imbalanced: *Streptococcus pneumoniae* has 174 genomes, *E. coli* 97, and *Klebsiella pneumoniae* 47, whereas 16 taxa have a single genome. PREMISE attaches each genome as a leaf below its taxon in the NCBI taxonomy, giving a tree of 473 vertices; the methods that take a taxonomy (Centrifuger and Ganon) were given the same NCBI taxonomy.

Table S1: **Benchmark datasets.** The 48 benchmark samples form six datasets. The dense dataset comprises the first ten CAMI II strain-madness samples [28], simulated from a pool of 408 genomes; for each sample, we report the number of genomes that contribute reads. Three sparse datasets consist of read pairs simulated by InSilicoSeq [20] in three scenarios: viral isolates, mixtures of three viruses of different subtypes, and mixtures of three strains of the same subtype; their rows correspond to the samples Isolate-1 to Isolate-10, Subtype-1 to Subtype-10, and Strain-1 to Strain-10. The two remaining sparse datasets consist of real samples. We report the subtype determined by the HA and NA segments (for subtype mixtures, in decreasing order of abundance), the strain identifiers of the strain mixtures, the SRA accessions of the real samples, the read length in base pairs (bp), and the number of read pairs, in thousands, before (**Orig.**) and after (**Filt.**) the filtering that defines the ground truth, with the retained percentage in parentheses.

| **Dataset** | | | **Read** | | |
|---|---|---|---|---|---|
| | | | **Length** | **Counts** (×1000) | |
| **Source** | **Type** | **Details** | **(bp)** | **Orig.** | **Filt. (%)** |
| CAMI II | Strain madness | Sample 0 (405 genomes) | 150 | 6,655 | 6,655 |
| | | Sample 1 (406 genomes) | 150 | 6,656 | 6,656 |
| | | Sample 2 (406 genomes) | 150 | 6,656 | 6,656 |
| | | Sample 3 (408 genomes) | 150 | 6,654 | 6,654 |
| | | Sample 4 (407 genomes) | 150 | 6,654 | 6,654 |
| | | Sample 5 (408 genomes) | 150 | 6,651 | 6,651 |
| | | Sample 6 (405 genomes) | 150 | 6,654 | 6,654 |
| | | Sample 7 (406 genomes) | 150 | 6,655 | 6,655 |
| | | Sample 8 (406 genomes) | 150 | 6,657 | 6,657 |
| | | Sample 9 (407 genomes) | 150 | 6,655 | 6,655 |
| Synthetic | Isolate | H3N2 | 301 | 500 | 500 |
| | | H5N8 | 301 | 500 | 500 |
| | | H6N8 | 301 | 500 | 500 |
| | | H5N3 | 301 | 500 | 500 |
| | | H1N8 | 301 | 500 | 500 |
| | | H13N3 | 301 | 500 | 500 |
| | | H7N9 | 301 | 500 | 500 |
| | | H1N2 | 301 | 500 | 500 |
| | | H1N2 | 301 | 500 | 500 |
| | | H1N1 | 301 | 500 | 500 |
| Synthetic | Subtype mixture | H7N1, H3N9, H7N1 | 301 | 500 | 500 |
| | | H1N1, H5N7, H9N7 | 301 | 500 | 500 |
| | | H1N1, H9N2, H5N5 | 301 | 500 | 500 |
| | | H5N9, H1N2, H5N2 | 301 | 500 | 500 |
| | | H1N2, H3N1, H5N3 | 301 | 500 | 500 |
| | | H7N1, H5N2, H9N2 | 301 | 500 | 500 |
| | | H1N8, H1N6, H7N1 | 301 | 500 | 500 |
| | | H3N2, H9N2, H9N1 | 301 | 500 | 500 |
| | | H1N8, H3N8, H9N1 | 301 | 500 | 500 |
| | | H7N3, H9N1, H1N2 | 301 | 500 | 500 |
| Synthetic | Strain mixture | H9N2 (C2203, C3109, C4329) | 301 | 500 | 500 |
| | | H1N1 (P14, P15, P18) | 301 | 500 | 500 |
| | | H3N2 (CY009, FY028, CY053) | 301 | 500 | 500 |
| | | H5N2 (a158, a504, a556) | 301 | 500 | 500 |
| | | H5N8 (SH-9, QD5, WFSG1) | 301 | 500 | 500 |
| | | H7N9 (PD-01, PD-02, Shanghai/01) | 301 | 500 | 500 |
| | | H7N8 (16-001403-1, 16-001573-2, AH0012935) | 301 | 500 | 500 |
| | | H5N1 (3472, 4571, 14756) | 301 | 500 | 500 |
| | | H3N2 (12TOSU268, A01259003, Indiana/21) | 301 | 500 | 500 |
| | | H1N1 (HL-1125, Orenburg/01, Giessen/6) | 301 | 500 | 500 |
| Real | Isolate | SRR31013463 | 301 | 666 | 643 (96.5) |
| | | SRR31013465 | 301 | 562 | 545 (96.9) |
| | | SRR31013467 | 301 | 495 | 426 (86.0) |
| | | SRR31013473 | 301 | 451 | 439 (97.5) |
| Real | Mixture | SRR3360139 | 251 | 1,202 | 1,140 (94.8) |
| | | SRR3360140 | 251 | 826 | 790 (95.6) |
| | | SRR3360145 | 251 | 1,071 | 1,023 (95.6) |
| | | SRR3360146 | 251 | 1,256 | 1,200 (95.5) |

*Ground truth.* CAMI records the genome from which each read was simulated. This genome is the true source of the read and, because CAMI genomes are strains, also its true label at the strain rank. The true label of a read at each rank from kingdom to species is the ancestor of the taxon of its genome at that rank. Six of the 26 taxa have no ancestor at the species rank, three have none at the genus rank, and one has none at the family rank; reads from their genomes have no true label at those ranks and are excluded from the evaluation at that rank (Section S5.3).

#### S5.2 Sparse Samples

In a sparse sample, a few of many highly homologous references are present: the 8 to 24 segments of one to three influenza viruses, drawn from a database of 5,268 segments. The challenge is to reject the many near-identical references that contribute no reads. We compared the methods in five scenarios, three synthetic with ten samples each and two real with four samples each (Table S1); the reference database, the synthetic and real samples, and the decoy references are described below.

*Influenza reference database.* The reference database for the influenza analyses was built from all Influenza A virus (IAV) sequences with complete assemblies in NCBI Virus [6] as of August 8, 2026. We retained only the latest version of each sequence and removed sequences with more than 20% ambiguous nucleotides, leaving 6,528 sequences. We then removed exact duplicates and sequences contained in a longer sequence, giving a final reference set of 5,268 IAV segments. The synthetic samples were drawn from this database, which also contains every strain present in the real samples.

*Synthetic samples.* The three synthetic datasets were generated with InSilicoSeq v2.0.1 [20]. Each synthetic virus was assembled by drawing, without replacement, one sequence of each segment from the reference database. Each sample consists of 500,000 read pairs of 301 bp, simulated with the built-in MiSeq error profile from the eight segments of one virus for the isolates and from the 24 segments of three viruses for the subtype and strain mixtures. The curation scripts, random seeds, and configuration needed to reproduce the database and the synthetic samples are deposited in Zenodo (`https://doi.org/10.5281/zenodo.19026876`). For the synthetic samples, the true source of every read and the true abundances are known.

*Real samples.* The first real dataset consists of total RNA sequencing data from four avian influenza isolates, generated by the Agricultural Research Service, United States Department of Agriculture, and described by Goraichuk et al. [13]. The samples were sequenced on an Illumina MiSeq ($2 \times 301$ bp read pairs) following standard IAV processing practices [31]. The reads had been quality trimmed and filtered before deposition, as described by Goraichuk et al. [13], but SISPA [9] primer sequences remained in some reads. We trimmed these primers with cutadapt v5.2 [26], using the options `-g GACCATCTAGCGACCTCCACNNNNNNNN`, `-G GACCATCTAGCGACCTCCACNNNNNNNN`, `--overlap 8`, `--minimum-length 20`, and `--trim-n`. The trimmed reads were aligned to their respective references (eight segments) with BWA-MEM2 v2.2.1 [43]. Read pairs aligned to the same reference segment, with no supplementary (split) alignment for either read, were retained for analysis. The references of these alignments were taken as the true classifications, and the true relative abundance $\pi_l$ of each of the eight reference segments $l$ was computed as the number of read pairs assigned to $l$ divided by the number of retained read pairs.

The second real dataset comprises four paired-end viral RNA whole-genome sequencing datasets from the NCBI Sequence Read Archive (SRA), BioProject PRJNA317621 [27]. These samples are defined two-strain mixtures of the PR8 and WSN33 IAV (H1N1) strains at known ratios, with the minor strain making up between 5% and 0.63% of the mixture. Unlike the isolate reads, these reads carried no SISPA primers, so we used cutadapt v5.2 [26] only to remove the 3′ Illumina read-through adapter and to discard reads shorter than 30 bp, with the options `-a AGATCGGAAGAGC`, `-A AGATCGGAAGAGC`, and `--minimum-length 30`. The trimmed reads were aligned with BWA-MEM2 v2.2.1 [43] to a reference of the two known strain genomes (16 segments, `PR8_<SEG>` and `WSN33_<SEG>`). Read pairs aligned to the same reference segment, with no supplementary (split) alignment for either read, were retained, as for the isolates, and their BWA-MEM2 references define the ground truth.

Real samples contain contaminants, chimeras, and other reads without a true source in the reference, so this ground truth is an estimate. We therefore estiamted $\widehat{\boldsymbol{\pi}}$ over the BWA-classified reads only and report coverage instead of recall for the real samples (Section S5.3).

*Decoy references.* In the original database, the closest absent reference to each true segment of the real samples is far less similar than the strains that co-occur in the synthetic strain mixtures. We therefore added near-identical decoys of every true segment, so that the real samples, like the synthetic ones, test the rejection of absent references. For each of the 48 true segments of the eight real samples (32 from the isolates and 16 from the PR8/WSN33 mixtures), we created two decoys, 96 in total, by substituting the nucleotides at $m$ positions chosen uniformly at random without replacement, 60% of the substitutions being transitions. The target similarity of each decoy to its source was drawn from the 31-mer containment similarities between co-occurring strains of the same segment in the synthetic strain mixtures (median 0.94, range 0.80–0.98). The number of substitutions $m$ was set so that the expected containment matched the target, and the positions were redrawn until the achieved containment was within 0.01 of the target. The two decoys of a segment share half of their substitutions, so the source and its two decoys are about equally similar to one another. The decoys are part of the reference given to every method for all influenza samples, but only the real samples contain the nearby sources.

#### S5.3 Evaluation Metrics

We evaluated the methods on the precision and coverage of their read classifications, their false positive (FP) and false negative (FN) reference identifications, the accuracy of their abundance estimates $\widehat{\boldsymbol{\pi}}$, and their runtime and memory usage [47]. The methods differ in what they report: PREMISE, Centrifuger, Ganon, KMCP, Mora, and Salmon classify individual reads and report a profile, whereas Karp and Sylph report only a profile and are therefore scored on the reference- and abundance-level metrics alone.

At the read level, precision is the proportion of classified reads that are classified correctly, and coverage is the proportion of input reads that are classified. Recall is the proportion of input reads that are classified correctly, and the F1 score is the harmonic mean of precision and recall. For the real samples, which may include artefactual reads, we report coverage instead of recall.

At the reference level, let $\mathcal{S}$ be the set of references present in a sample, i.e., with true abundance $\pi_s > 0$, and let $\widehat{\mathcal{S}}$ be the set of references that a method identifies, i.e., with estimated abundance $\widehat{\pi}_s > 0$. The Sørensen similarity, $2|\mathcal{S} \cap \widehat{\mathcal{S}}|/(|\mathcal{S}| + |\widehat{\mathcal{S}}|)$, measures the accuracy of source detection, which is critical for diagnostic validity; the ablation study (Section S8) reports the closely related Jaccard similarity, $|\mathcal{S} \cap \widehat{\mathcal{S}}|/|\mathcal{S} \cup \widehat{\mathcal{S}}|$. The numbers of FP and FN references are $|\widehat{\mathcal{S}} \setminus \mathcal{S}|$ and $|\mathcal{S} \setminus \widehat{\mathcal{S}}|$, respectively.

The accuracy of the abundance estimates is measured by the Ruzicka similarity between $\widehat{\boldsymbol{\pi}}$ and $\boldsymbol{\pi}$. The Ruzicka similarity suits compositional data: it penalizes discrepancies in relative abundance, and one minus it satisfies the properties of a metric [44]. For the methods that estimate abundances natively (PREMISE, Karp, KMCP, Salmon, and Sylph), $\widehat{\boldsymbol{\pi}}$ is taken directly from their output. For Centrifuger, Ganon, and Mora, $\widehat{\boldsymbol{\pi}}$ is computed from the number of reads assigned to each reference, divided by the total number of assigned reads, so unclassified reads are excluded.

On the dense samples, every metric is computed at each taxonomic rank $R$ from kingdom to strain, with the truth and the predictions both replaced by their ancestors at $R$, or at the strain rank, where the truth and the predictions are genomes. A read classified to a taxon coarser than $R$ counts as unclassified at $R$; examples are a read that PREMISE stops above the species rank, or above the genomes of its species, and a read that Centrifuger places at a lowest common ancestor. Reads whose true taxon has no ancestor at $R$ (Section S5.1) are excluded from the read-level metrics at $R$, and the abundance of the taxa without an ancestor at $R$ is removed from both the true and the predicted profiles, which are then renormalized. At the strain rank, every read has a true genome, so no read is excluded, and the reference-level metrics count genomes: a genome is detected if its estimated abundance is positive.

### S6 Index Construction

To compare construction time and index size, we built an index from each of the influenza and CAMI II reference sets with every method. All methods used the same reference set and their default parameters; PREMISE used all available cores, Ganon, KMCP, Salmon, and Sylph used 40 threads, and the remaining methods used their default number of threads. Table S2 summarizes the results; the times and sizes exclude the taxonomy that Centrifuger downloads and stores.

On the influenza reference, PREMISE builds a 45.25 MB B-RLFM in 1.57 s. Sylph builds the smallest index, and builds it fastest (1.97 MB in 0.04 s), followed in size by KMCP (3.56 MB) and Ganon (6.60 MB), whose build takes much longer (25.01 s). Centrifuger builds the next smallest index (21.63 MB in 6.33 s), and Mora, Karp, and Salmon build indexes of 50.95 MB, 76.19 MB, and 89.22 MB in 2.23 s, 3.66 s, and 1.04 s, respectively. The CAMI II reference (1.45 Gbp) is about 160 times larger than the influenza reference (9.1 Mbp), but the B-RLFM of PREMISE grows only 40-fold, to 1.83 GB; the indexes of Mora (8.00 GB), Karp (4.38 GB), and Salmon (2.17 GB) are larger, and those of Ganon, KMCP, Centrifuger, and Sylph are smaller. PREMISE builds it in 253 s, faster than Mora (307 s), Karp (605 s), and Centrifuger (4,503 s), but slower than Salmon (217 s) and the $k$-mer-based Ganon, KMCP, and Sylph (under 15 s). On this reference too, Sylph builds the smallest index, and builds it fastest (0.11 GB in 0.36 s), followed in size by Centrifuger (0.33 GB), which has the longest build time (4,503 s). Sampling its suffix array makes the B-RLFM several times smaller (Section S8).

Table S2: **Index construction.** Each index is built from the same reference collection; size is that of the final index required for classification. (a) The influenza reference, with size in MB ($10^6$ bytes); (b) the CAMI II strain-madness reference, with size in GB ($10^9$ bytes). Build time is in seconds.

(a) Influenza

| Method | Size (MB) | Time (s) |
|---|---|---|
| PREMISE | 45.25 | 1.566 |
| Centrifuger | 21.63 | 6.326 |
| Ganon | 6.601 | 25.01 |
| Karp | 76.19 | 3.660 |
| KMCP | 3.559 | 0.8074 |
| Mora | 50.95 | 2.228 |
| Salmon | 89.22 | 1.041 |
| Sylph | 1.973 | 0.03996 |

(b) CAMI II

| Method | Size (GB) | Time (s) |
|---|---|---|
| PREMISE | 1.829 | 252.7 |
| Centrifuger | 0.3274 | 4503 |
| Ganon | 0.8460 | 14.62 |
| Karp | 4.383 | 605.2 |
| KMCP | 0.5142 | 12.17 |
| Mora | 8.003 | 307.1 |
| Salmon | 2.169 | 217.1 |
| Sylph | 0.1143 | 0.3608 |

### S7 Comparative Results

This section reports the per-method and per-scenario results, together with the false and missed taxa on CAMI II (Figure S1) and the runtime and memory of every method (Figure S2 and Figure S3).

#### S7.1 CAMI II

On the ten CAMI II strain-madness samples, PREMISE was run over the NCBI taxonomy without the sparsity penalty ($\rho = 0$) for a fixed 2,000 EM iterations and assigned reads at $\gamma = 0.5$; all eight methods completed every sample. Every method was scored at each rank from kingdom to species and at the strain rank (Section S5.3).

*Classification and profiling across ranks.* From kingdom to family, every method that classifies reads assigns nearly all of them correctly (precision $\geq$ 0.999 and coverage $\geq$ 0.998, except for the coverage of KMCP, 0.988), whereas the abundance profiles of KMCP and Karp already deviate at the phylum rank (Ruzicka similarity 0.680 and 0.669). The methods separate at the genus and species ranks. At the species rank, PREMISE has the highest precision (0.9986; 0.9967–0.9994) and coverage (98.72%), and hence the highest F1 score (0.992), ahead of Ganon (0.980), Mora (0.971), KMCP (0.959), Salmon (0.956), and Centrifuger (0.879), whose coverage falls to 78.98% because it places reads with tied hits at their lowest common ancestor. PREMISE, Centrifuger, Ganon, Mora, and Salmon detect exactly the species present (Sørensen similarity 1.0; no FP or FN at any rank down to the species), while KMCP and Sylph miss 1.9 and 3.1 species per sample on average. Salmon estimates species abundances most accurately (Ruzicka similarity 0.978), followed by PREMISE (0.966) and Mora (0.955); Ganon (0.909), Sylph (0.881), Centrifuger (0.756), KMCP (0.660), and Karp (0.252) trail. The difference between PREMISE and Salmon lies in one pair of taxa: PREMISE assigns 2–5% of the abundance of each sample from genomes labeled only with the family Enterobacteriaceae to *E. coli* genomes, with which they share 17% of their sequence verbatim (17 Mbp).

*Strain-level classification and profiling.* At the strain rank, each read must be traced to the genome from which it was simulated, among up to 174 genomes of the same species. Ganon, Mora, Salmon, and KMCP assign nearly every read to a genome, but only 32.6%, 18.1%, 15.9%, and 15.5% of the assignments are correct, so their precision equals their recall. PREMISE has the highest strain precision (0.804; 0.732–0.862) and F1 score (0.493, against 0.326 for Ganon), assigning 44.8% of the reads to a genome and the rest to their species. Centrifuger, which places reads with tied hits at their lowest common ancestor, assigns 15.0% of the reads to a genome at a precision of 0.639. PREMISE and Salmon estimate genome abundances far more accurately than the other methods (Ruzicka similarity 0.681 and 0.680, against at most 0.283 for Mora), each ahead on five of the ten samples. None of PREMISE, Centrifuger, and Mora misses a genome present

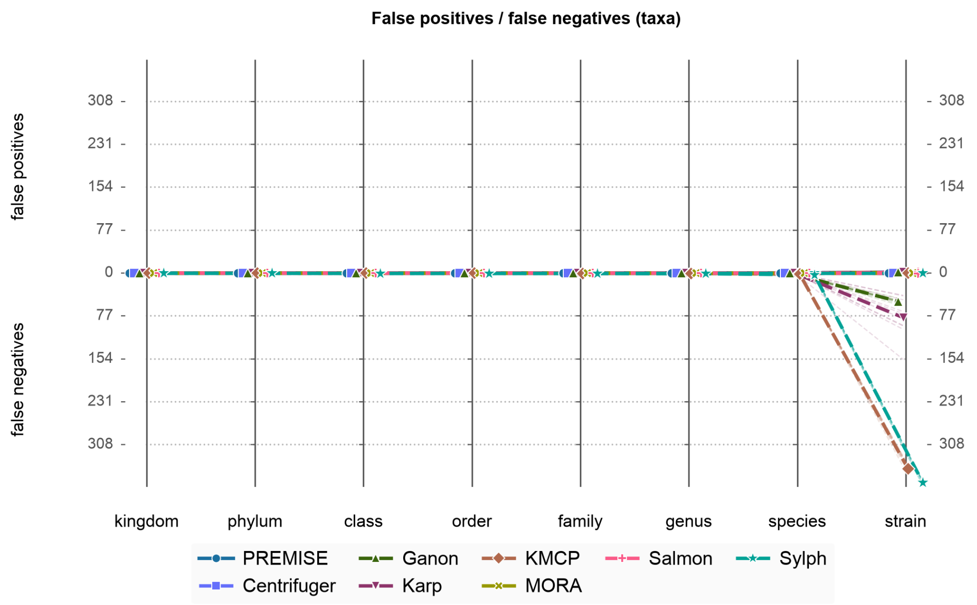


Fig. S1: **False and missed taxa across taxonomic ranks on CAMI II.** Mean number of false positive (FP, above the horizontal axis) and false negative (FN, below) taxa per sample at every rank from kingdom to species and genomes at the strain rank; faint lines are the ten samples. Every method detects exactly the taxa present down to the genus; KMCP and Sylph miss species, and at the strain rank, Ganon, Karp, KMCP, and Sylph miss genomes present in the sample.

in a sample, and Salmon misses 0.8 per sample, whereas Ganon, Karp, KMCP, and Sylph miss on average 51, 80, 351, and 376 of the 405 to 408 genomes present (Figure S1). PREMISE, Centrifuger, Karp, Mora, and Salmon also report the 0 to 3 reference genomes absent from each sample (1.6 on average).

*Resource utilization.* Every method completed each CAMI II sample within about 20 minutes (Figure S2). Sylph is the fastest (10.4 s, 1.0 GB), and Centrifuger is the most memory-efficient (106 s, 0.42 GB). The search of KMCP takes 283 s with 1.84 GB, and its profiling step another 6.5–15 minutes per sample. Ganon (317 s, 22.4 GB) and Mora (354 s for the BWA-MEM2 alignment, plus 33–34 s for Mora itself) take similar times, followed by Salmon (531 s, 19.5 GB), PREMISE (572 s; 546–594 s), and Karp (845 s, 12.2 GB). PREMISE needs more memory than every method except Mora (41.8–49.5 GB, 45.9 GB on average, against 52.9 GB for Mora).

#### S7.2 Influenza

On the influenza samples, PREMISE was run without a taxonomy and with the sparsity penalty ($\rho = 150$, $\omega = 10^{-10}$) for a fixed 2,000 EM iterations, and assigned reads at $\gamma = 0.6$. For each scenario, we report the read-level precision, coverage, and F1 score, the reference-level FP, FN, and Sørensen similarity, and the abundance accuracy (Ruzicka similarity), with samples labeled as in Table S1.

*Synthetic samples.* On the synthetic isolates, PREMISE recovers exactly the true references of all ten samples, with no false or missed references (Sørensen similarity 1.0), perfect precision (1.0), coverage of 99.44%–99.86%, and Ruzicka similarity $\geq 0.992$. Among the other methods, Centrifuger achieves high precision (0.917–0.983) but lower coverage (72.05%–91.72%). Salmon and Ganon assign 100.00% of the reads, but Salmon reports many false references (138–236; precision 0.602–0.899), and Ganon has lower precision (0.532–0.907) on all isolates but Isolate-3. Centrifuger, Karp, and Mora miss no references but report many false ones (up to 1,321 for Karp), whereas Ganon misses up to three present references and Sylph misses three to six of the eight in every sample (Ruzicka similarity 0.083–0.302). PREMISE has the highest Ruzicka similarity on every isolate except Isolate-3, where the profile of Ganon is exact.

On the subtype mixtures, Karp, Centrifuger, and Mora report hundreds to thousands of false references (up to 1,784 for Karp). PREMISE recovers exactly the true references of all ten samples, with no false

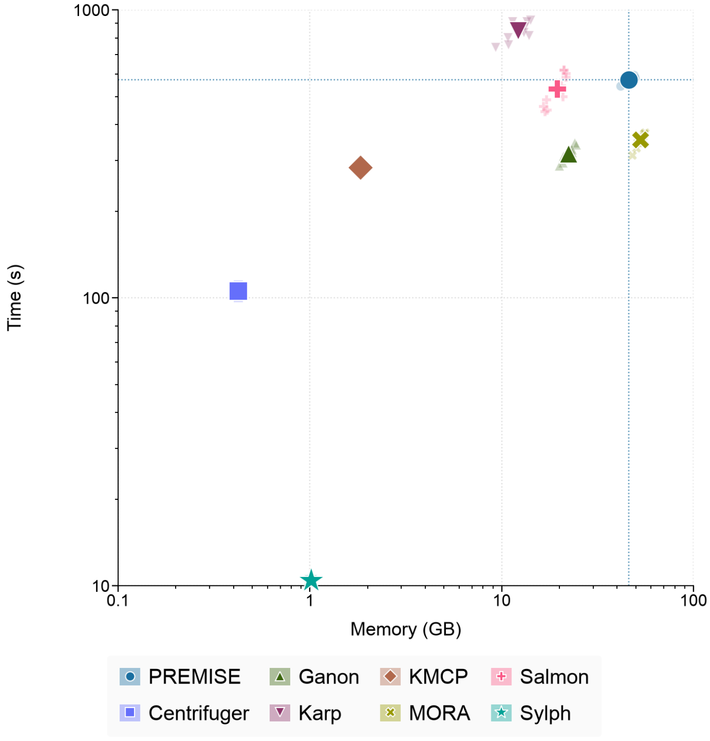


Fig. S2: **Resource utilization on CAMI II.** Log-scaled run-time in seconds ($y$-axis) against log-scaled peak memory in GB ($x$-axis) required by each method to analyze a single CAMI II sample (6.65 million read pairs). Faint points are individual samples and large points the mean of each method; the dotted lines cross at the mean of PREMISE. The times of KMCP and Mora cover `kmcp search` and the BWA-MEM2 alignment only, excluding `kmcp profile` and the Mora reassignment step.

or missed references (Sørensen similarity 1.0), a read precision of 0.997–1.000, coverage of 98.33%–99.80%, and Ruzicka similarity $\geq$ 0.995. Salmon assigns every read (100.00%), misses no references (FN = 0), and estimates abundances well (Ruzicka similarity 0.861–0.983, below PREMISE on every sample), but it reports 336–412 false references. Ganon has the next best Sørensen similarities (0.436–0.784), with up to 37 false and 8 missed references, whereas the read coverage of Centrifuger drops to 67.30% and Sylph misses 12–17 of the 24 present references in every sample.

The strain mixtures test the resolution of the methods, since their strains share most of their sequence. Precision drops sharply for Ganon (0.163–0.824) and Mora (0.275–0.942), and the coverage of Centrifuger falls to 24.85% on Strain-1. Salmon assigns every read (100.00%), recovers all sources (FN = 0), and estimates abundances well (Ruzicka similarity 0.878–0.993), but it reports up to 457 false references. PREMISE keeps a precision of 0.849–0.998 and recovers exactly the true references of eight of the ten samples; it reports two false references and misses one on Strain-1 and misses one on Strain-6, but its read coverage varies widely (55.20%–99.26%, mean 77.77%). Centrifuger, Mora, and Karp report up to 701, 687, and 1,239 false references, respectively, while Sylph misses between 15 and all 24 true references per sample.

Across all synthetic samples, PREMISE has the highest read-level precision, except on Strain-1, Strain-5, Strain-6, and Strain-7, where Centrifuger, classifying 24.85%, 86.30%, 73.91%, and 55.98% of the reads, is more precise. PREMISE and Salmon are the two most accurate abundance estimators: PREMISE has the highest Ruzicka similarity on nine, ten, and eight of the ten isolates, subtype mixtures, and strain mixtures, Salmon on the other two strain mixtures (Strain-1 and Strain-5), and Ganon on Isolate-3. On the reads that Centrifuger classifies, PREMISE is more precise than Centrifuger on nine of the ten strain mixtures (Barnard's exact test [3], one-sided $p < 10^{-6}$; $p = 0.96$ on Strain-3).

*Real samples.* The reference of both real scenarios contains the near-identical decoys of every true segment (Section S5.2), and many of the false references below are decoys: PREMISE detects one decoy in one isolate and none in the other isolates or the mixtures, whereas Ganon detects 7–12 per sample, Centrifuger 16–30, Salmon 24–33, Mora 26–33, and Karp 44–76. On the isolates, PREMISE maintains near-perfect read-level precision (1.00 on three isolates and 0.998 on SRR31013465) while assigning 96.7%–97.9% of the reads; it recovers exactly the eight segments of three isolates, reports two false references on SRR31013465 (a decoy and one other reference), misses no reference, and reaches a mean Ruzicka similarity of 0.995 (0.991–0.998), the highest of all methods on every isolate. Centrifuger keeps high precision (mean 0.89, range 0.74–0.99) but assigns only 21%–43% of the reads (mean 32%) and reports 74–340 false references per sample (mean 192; mean Ruzicka similarity 0.46). Ganon assigns 99.7% of the reads at a lower precision (mean 0.82, range 0.66–0.92), with 65–142 false references (mean 101) and a mean Ruzicka similarity of 0.71, the best among the other methods. On the three isolates that it completed, KMCP reports no false references but misses four to six of the eight segments, at a precision of 0.49–0.68.

On the mixtures, PREMISE recovers the 16 true segments of both strains in every sample with no false or missed references (Sørensen similarity 1.0), at a read-level precision of 1.0, with 99.5%–99.6% of the reads assigned and a mean Ruzicka similarity of 0.997. Centrifuger keeps a high read precision (0.99) but assigns only 22.6%–23.7% of the reads, reports 45–70 false references per sample (mean 60), misses one true segment in three of the four samples (mean Sørensen similarity 0.34), and estimates abundances poorly (mean Ruzicka similarity 0.54). Ganon assigns nearly all reads, but at a precision of only 0.80–0.81; it reports 33–80 false references per sample (mean 58), misses one to three true segments per sample (mean Sørensen similarity 0.33), and estimates abundances with substantial error (mean Ruzicka similarity 0.67). Karp reports 2,416–2,851 false references per sample (mean 2,620), with a mean Ruzicka similarity of 0.38 and Sørensen similarity of 0.01. KMCP assigns 99.7% of the reads at a precision of only 0.52–0.54, but its profiling step discards all references and returns an empty profile for every sample. Mora assigns 97.6%–97.9% of the reads at a precision of 0.58–0.59 and reports 56–68 false references per sample (mean 61), with a mean Ruzicka similarity of 0.41 and Sørensen similarity of 0.35. Salmon assigns 99.9% of the reads at a precision of 0.84–0.85; it produces the next best abundance estimates (mean Ruzicka similarity 0.98) and reports 52–62 false references per sample (mean 56; mean Sørensen similarity 0.36). Sylph reports 11–16 false references while missing six to eight of the 16 true segments per sample, with a mean Ruzicka similarity of 0.25 and Sørensen similarity of 0.45.

*Resource utilization.* Across the influenza samples, the methods fall into two groups by their computational requirements, with KMCP as an outlier (Figure S3). The methods based on sketches or $k$-mer lookups are the fastest: Sylph completes in under a second, Centrifuger within 5 s, and Ganon and Karp within 36 s, all using less than 1.8 GB of memory. The alignment- and likelihood-based methods need more resources: PREMISE (13.1–39.9 s, 0.73–2.53 GB) and Salmon (7.0–49.3 s, 1.24–7.69 GB) remain modest, whereas Mora needs up to 13.8 GB. PREMISE uses 4–7× less memory than Mora in every scenario, and its mean runtime is below those of Salmon and Mora on the synthetic samples (28–29 s); on the real samples, Mora is faster on the isolates (13.2 s vs. 16.4 s), and Salmon on the mixtures (10.9 s vs. 30.8 s, with Mora at 30.5 s). The runtime of Salmon varies more (7–36 s on the real isolates vs. 9–12 s on the real mixtures). KMCP is a slow outlier on the real isolates (11–34 min per sample) and did not finish SRR31013465 within the one-hour limit, although it finished each real mixture in under a minute; it also did not finish any synthetic sample within the limit, where we estimate that `kmcp search` alone would take about 3.6 h per sample. Overall, PREMISE needs about an order of magnitude more time and memory than the sketching tools on the influenza samples.

### S8 Ablation Study

To evaluate the stability of PREMISE, we performed a one-at-a-time sensitivity analysis, sweeping each of six parameters while holding the others at their defaults: the minimum seed length $\kappa$ (`-m`, default 22), the likelihood $\epsilon_1$ below which a read is left unclassified (default 0), the relative likelihood $\epsilon_2$ below which a candidate reference is dropped for a read (default $10^{-18}$), the penalty weight $\rho$ (default 150), the penalty scale $\omega$ (default $10^{-10}$), and the sampling rate of the suffix array in the B-RLFM (default 1, the full suffix array). The sweeps used the four real isolates and the four real mixtures, with the sparse-sample settings of Section S7.2, including the fixed 2,000 EM iterations. Figures S4 and S5 show how the first five parameters affect precision, coverage, Ruzicka similarity, Jaccard similarity, and runtime. Below, values are means over

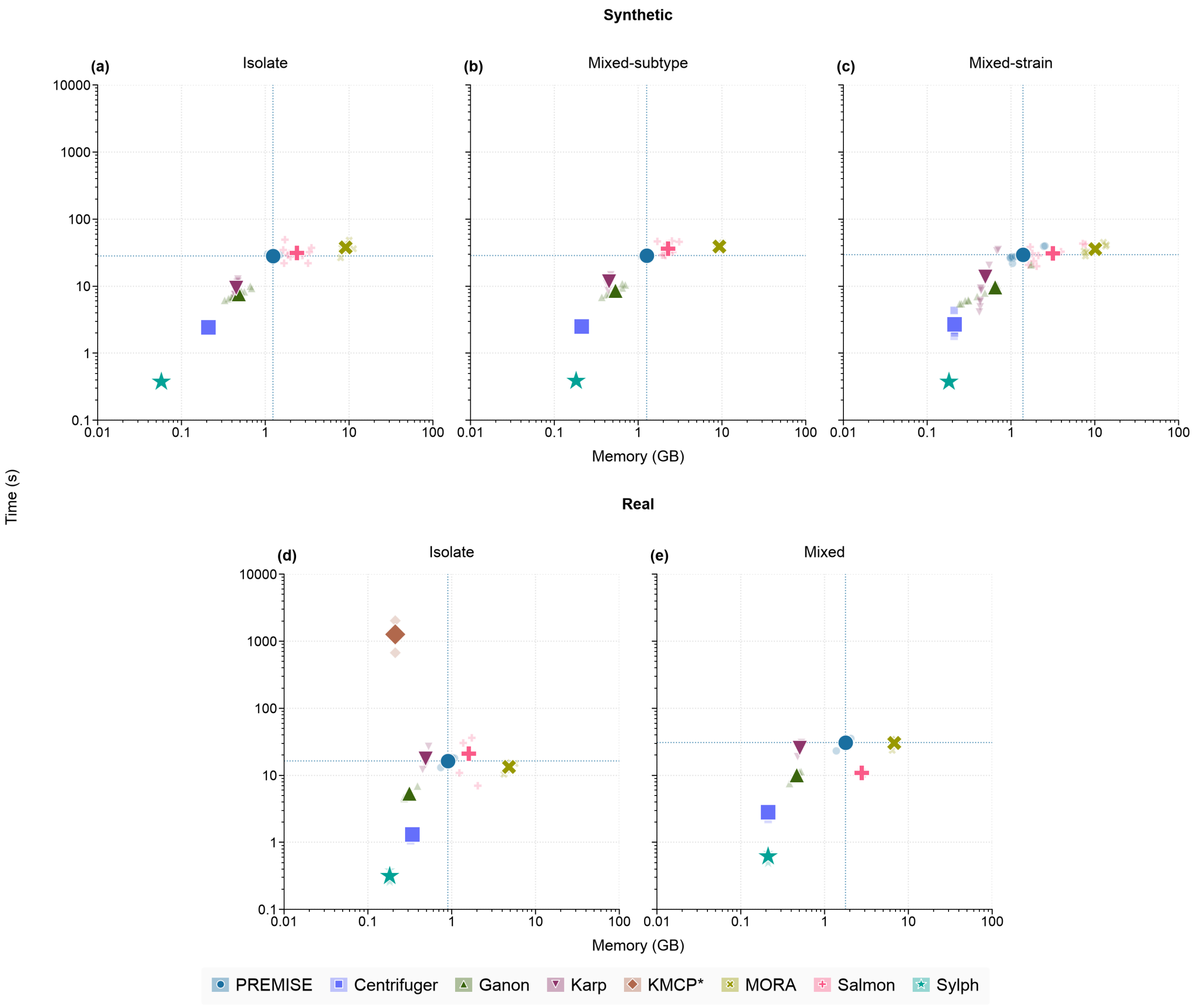


Fig. S3: **Resource utilization on the influenza samples.** Each scatter plot shows the log-scaled runtime in seconds ($y$-axis) against the peak memory in GB ($x$-axis) required by each method to analyze a single sample. The top row (a–c) shows the synthetic isolates, subtype mixtures, and strain mixtures, and the bottom row (d–e) the real isolates and mixtures. Faint points are individual samples and large points the mean of each method; the dotted lines cross at the mean of PREMISE. KMCP (asterisk) did not finish any synthetic sample or one real isolate within the one-hour limit.

the four samples of a scenario, and abundance accuracy and source detection are reported as the Ruzicka and Jaccard similarities (Section S5.3).

*Seed length.* Varying the minimum SMEM length $\kappa$ from 20 to 24 leaves precision and Jaccard similarity unchanged in both scenarios (0.9995 and 0.932 on the real isolates, and 1.0 and 1.0 on the real mixtures). Longer seeds drop a few distant matches, so coverage falls slightly, from 97.58% to 97.36% on the real isolates and from 99.60% to 99.55% on the real mixtures; the Ruzicka similarity changes by less than 0.0003. The mean runtime falls steadily with longer seeds, from 15.7 to 14.4 s per real isolate and from 27.8 to 26.7 s per real mixture, and the peak memory stays at about 0.90 GB per real isolate and 1.78 GB per real mixture. We use $\kappa = 22$ by default.

*Alignment filtering ($\epsilon_1$, $\epsilon_2$).* Filtering candidate alignments before EM has little effect on accuracy over broad ranges of both thresholds. Raising the relative threshold $\epsilon_2$ from $10^{-36}$ to $10^{-9}$ leaves every accuracy measure, the runtime, and the peak memory essentially unchanged in both scenarios. Raising the absolute

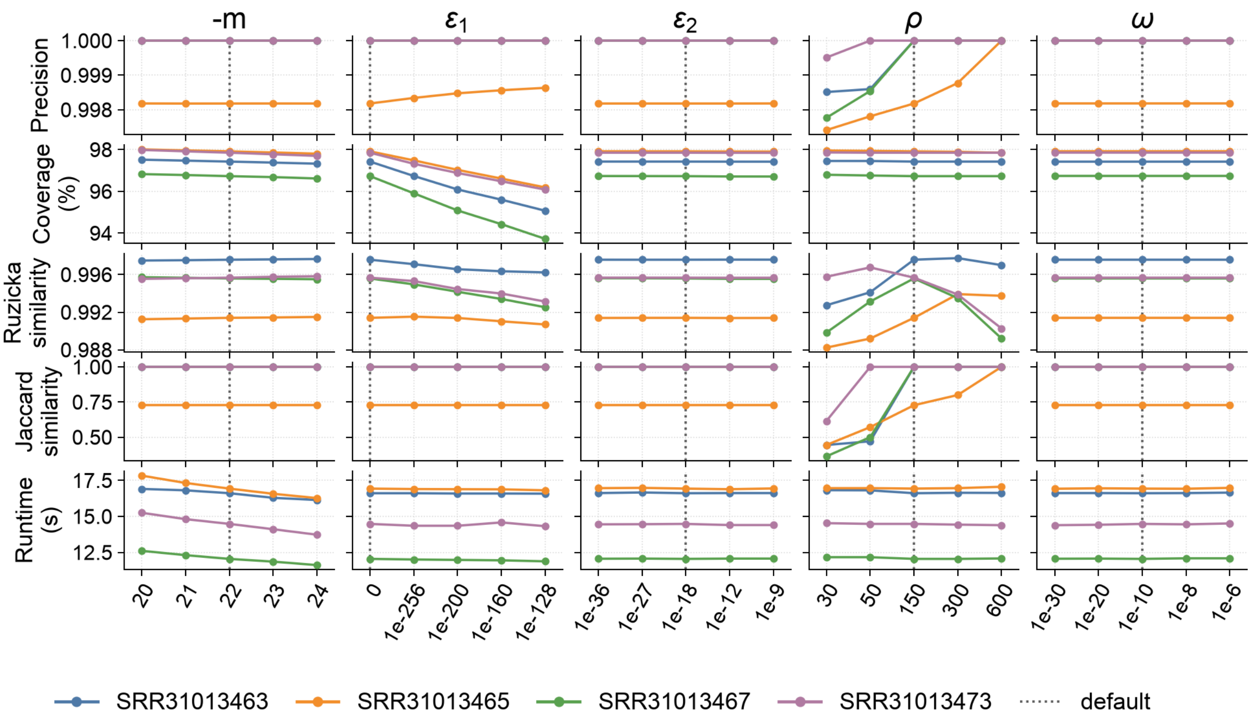


Fig. S4: **Ablation study on the real isolates.** Each column varies one parameter of PREMISE (left to right: the minimum seed length `-m`, $\epsilon_1$, $\epsilon_2$, $\rho$, and $\omega$), and each row shows one measure (top to bottom: precision, coverage, Ruzicka similarity, Jaccard similarity, and runtime) on the four real isolates; each line is one sample, and the dotted line marks the default value.

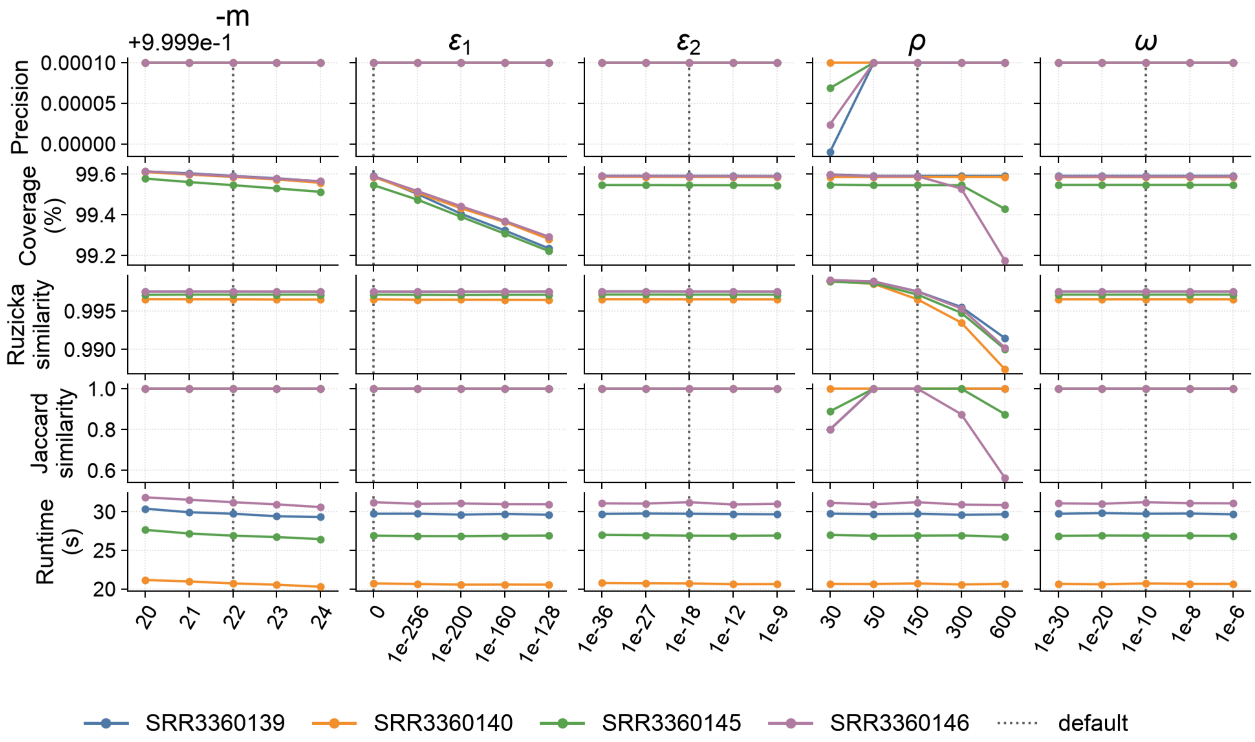


Fig. S5: **Ablation study on the real mixtures.** Each column varies one parameter of PREMISE (left to right: the minimum seed length `-m`, $\epsilon_1$, $\epsilon_2$, $\rho$, and $\omega$), and each row shows one measure (top to bottom: precision, coverage, Ruzicka similarity, Jaccard similarity, and runtime) on the four real PR8/WSN33 mixtures; each line is one sample, and the dotted line marks the default value.

threshold $\epsilon_1$ from 0 to $10^{-128}$ acts directly on coverage, which falls from 97.48% to 95.27% on the real isolates and from 99.58% to 99.26% on the real mixtures. Precision rises slightly on the real isolates (from 0.99955 to 0.99966), Jaccard similarity is unchanged in both scenarios, and the Ruzicka similarity falls from 0.9951 to 0.9932 on the real isolates and by less than 0.0001 on the real mixtures. The defaults are $\epsilon_1 = 0$ and $\epsilon_2 = 10^{-18}$.

*Penalty weight and scale ($\rho$, $\omega$).* The penalty weight $\rho$ controls how many references EM prunes, and it is the only swept parameter that changes source detection. Weak penalties leave false references: at $\rho = 30$, the mean Jaccard similarity is 0.467 on the real isolates and 0.872 on the real mixtures, and at $\rho = 50$, it is 0.636 on the real isolates, although every real mixture is already exact. The default, $\rho = 150$, is the smallest swept value that also recovers exactly the segments of three of the four real isolates (per-sample Jaccard similarity 1.0, 0.73, 1.0, and 1.0), and it gives the most accurate abundances on the isolates (Ruzicka similarity 0.9951, against 0.9917 at $\rho = 30$, 0.9948 at $\rho = 300$, and 0.9926 at $\rho = 600$). The two false references left on SRR31013465 are pruned only by heavier penalties: its Jaccard similarity rises to 0.80 at $\rho = 300$ and to 1.0 at $\rho = 600$, where every isolate is exact and precision reaches 1.0. On the mixtures, however, heavier penalties shrink the proportions of the minor strains and eventually drop them: the Ruzicka similarity falls from 0.9972 at $\rho = 150$ to 0.9948 at $\rho = 300$ and 0.9898 at $\rho = 600$, and the Jaccard similarity falls to 0.969 and 0.859. Weaker penalties give more accurate abundances on the real mixtures (Ruzicka similarity 0.9989 at $\rho = 30$ and 0.9987 at $\rho = 50$), at the cost of false references on the isolates and, at $\rho = 30$, on the mixtures. The default thus balances the isolates, which favor heavier penalties, against the mixtures, which lose true strains under them. Varying the penalty scale $\omega$ across 24 orders of magnitude, from $10^{-30}$ to $10^{-6}$, changes no accuracy measure in either scenario by more than $10^{-4}$.

*Suffix-array sampling.* Keeping only every $k$th suffix-array entry shrinks the B-RLFM but slows down the location of seed occurrences; it does not change which occurrences are found. On the influenza reference, sampling every 64th entry instead of every entry shrinks the B-RLFM from 45.25 MB to 9.35 MB (27.02, 17.90, 13.34, 11.06, and 9.92 MB at rates 2, 4, 8, 16, and 32), with build times of 1.5–1.6 s. The mean alignment runtime grows by at most 6% (from 8.2 to 8.6 s per real isolate and from 14.4 to 15.0 s per real mixture), and the peak memory falls slightly (from 0.86 to 0.83 GB per real isolate and from 1.70 to 1.67 GB per real mixture). We keep the full suffix array by default, which favors speed; a sampled suffix array suits larger reference databases, where the size of the index matters more.
